# Mechanistic modeling of bacterial translation initiation across growth conditions

**DOI:** 10.64898/2026.08.31.748211

**Authors:** Jiahui Qin, Andreas Kremling

## Abstract

Translation frequency in bacteria depends on how ribosomes, mRNAs, and initiation factors are allocated across growth conditions. Here, we developed a mechanistic ODE-based model of *Escherichia coli* translation that represents initiation, elongation, termination, and coupled auxiliary processes. Growth-dependent abundances were derived from physiological relationships and reprocessed omics data, and simulated outputs were compared with translation-frequency and active-ribosome references. The model predicts a continuous shift from complex-formation-limited toward ribosome-limited behavior as growth increases. This shift is characterized by a decline in free-ribosome abundance, whereas initiation-factor pools remain largely unbound and do not become depleted in parallel. Together with the implemented IF-dependent kinetic term, this preserved availability provides a model-internal route through which productive initiation can be maintained despite increasing ribosome utilization. Consistently, transcript-wide ribosome loading remains below its theoretical maximum, while COG-level simulations reveal distinct sector-specific translation-frequency trajectories. The study therefore provides a resource-allocation framework for interpreting how mRNA–ribosome interactions shape bacterial translation across growth conditions.

## 1 Introduction

Protein synthesis is one of the largest and most tightly regulated investments in a growing bacterial cell. In *Escherichia coli*, changes in nutrient quality and growth rate reshape the abundance of ribosomes, mRNA, tRNA, translation factors, metabolic enzymes, and other cellular components [34, 6, 10, 38, 33, 11]. A central question is therefore not only how much translational machinery is present, but also how the available mRNA, ribosomes, and auxiliary factors are converted into productive protein synthesis across growth conditions.

Classical bacterial growth-law theory has provided a powerful framework for this problem by linking growth rate to proteome allocation. In this view, the cellular proteome is partitioned into functional sectors whose fractions change systematically with the growth rate: ribosome-associated proteins increase during faster growth, whereas metabolic or nutrient-acquisition sectors compensate according to the limiting condition [39, 26, 4, 33, 40]. This framework has clarified many coarse-grained principles of bacterial physiology, including proteome-economic explanations of overflow metabolism, growth-strategy trade-offs, and proteome-allocation programs across industrially relevant and distantly related bacteria [4, 32, 44, 49]. However, translation is not a protein-only process. Ribosome biogenesis requires a large investment in rRNA; mRNA provides the templates on which ribosomes initiate and elongate; tRNA and translation factors determine how efficiently ribosomes progress through the translation cycle [6, 10, 11]. Thus, a proteome-sector description alone cannot fully explain how growth-dependent molecular allocation is converted into translation flux.

This distinction is especially important when comparing proteome allocation with whole-biomass resource allocation. A ribosome-related protein sector may increase within the proteome while the total protein fraction relative to cell dry weight changes at the same time [6, 10, 47]. Conversely, the rise in total RNA with growth rate is driven primarily by rRNA rather than by a uniform increase of all RNA species. Therefore, a mechanistic interpretation of translation requires a common physiological scale on which protein, rRNA, tRNA, mRNA, and remaining biomass components can be considered together. The RNA-to-protein mass ratio provides such a growth-rate-dependent coordinate because it links the expansion of RNA-rich ribosome-associated biomass to global cellular composition.

At the same time, bulk resource allocation does not by itself determine translation performance. Translation frequency, here defined as the average number of proteins synthesized per mRNA template per unit time, depends on the kinetic interaction between mRNA and ribosomes [13, 27, 2]. Recent work has emphasized that protein production can be limited by different regimes, including mRNA–ribosome complex formation, ribosome availability, or transcript availability [8]. These regimes suggest that the same amount of cellular resource can produce different translation outputs depending on how efficiently ribosomes are recruited, initiated, elongated, terminated, and recycled.

## 1 Introduction

Translation initiation is a particularly important point of control in bacteria. Initiation requires productive engagement between the 30S ribosomal subunit and mRNA, recruitment of initiator tRNA, joining of the 50S subunit, and coordinated action of initiation factors IF1, IF2, and IF3 [5, 1, 19]. These factors regulate both the kinetics and fidelity of initiation-complex assembly. Their availability can therefore influence how effectively mRNA–ribosome encounters progress toward elongation-ready ribosomes.

Here, we developed and analyzed an ODE-based translation simulator to connect growth-rate-dependent resource allocation with mechanistic translation behavior in bacterial cells (represented by *E. coli*). The model incorporates initiation, elongation, termination, ribosome recycling, tRNA charging, and protein turnover. Growth-rate-dependent concentrations of the modeled components were derived from physiological relationships and reprocessed proteomic and transcriptomic datasets [6, 38, 33, 29, 47, 37, 13, 9]. To connect omics allocation with translation demand, genes were grouped using Clusters of Orthologous Groups (COG) annotations and then aggregated into three broad functional classes [42, 16]. In this coarse-grained mapping, COG1 represents information-processing and gene-expression-related functions, COG2 represents metabolism-related functions, and COG3 represents maintenance and other cellular-process functions [38, 27]. There are two other COG classes called COGx and COG_pseudo_ which represent the groups of genes with unknown function and the most idle genes respectively. More details of COG classification are provided in *Supplementary information*.

In the present model, the contribution of initiation factors is represented by an effective IF-dependent kinetic term rather than by separately resolved factor-binding pathways. The analysis addresses three linked questions: first, how protein, RNA, and COG-level omics allocations define the resources available for translation; second, how translation frequency changes with growth rate at the global and functional-sector levels; and third, how mRNA–ribosome distributions and initiation-factor availability support a growth-dependent crossover between limiting translation behaviors.

Together, these questions define translation frequency as a systems-level outcome of both resource availability and kinetic redistribution. Accordingly, the analysis first reconstructs the growth-dependent physiological and omics input space derived from various experimental data, then compares global and COG-level translation-frequency trajectories, and finally examines how mRNA occupancy, ribosome utilization, and initiation-factor availability change across the simulated growth range. This organization allows the model to be used not only as an isolated kinetic scheme but also as a bridge between biomass-scale resource allocation and the molecular distribution of the translation machinery. In this way, the study provides a mechanistic framework for interpreting how growth-dependent cellular composition is converted into mRNA–ribosome interaction and protein-production capacity in *E. coli*.

## 2 Results

### 2.1 A growth-rate-dependent resource landscape defines the translation input space

Before exploiting the ODE-based translation simulator, we reconstructed a growth-rate-dependent physiological and omics landscape to define the cellular resources available for protein synthesis in *E. coli*. This reconstruction integrates three layers of information: literature-derived physiological scaling relationships [6, 3, 27], biomass-level macromolecular composition [12, 23, 14, 43, 6, 15, 46, 3, 47, 2], and reprocessed proteomic and transcriptomic COG fractions [38, 33, 48, 13, 37, 9, 2]. Because the source datasets were obtained from different studies and reported on different reference scales, all quantities were converted to common reference forms. Global macromolecular quantities are expressed relative to cell dry weight (*ϕ*^*c*^ in g g_CDW_^−1^), whereas proteomic and transcriptomic profiles were normalized within samples and aggregated into COG-based functional groups. Protein mass fractions (*ϕ*^*P*^ in g g_P_^−1^) were converted into molar fractions (*χ*^*P*^ in mol mol_P_^−1^) using peptide lengths. Transcript signals were converted into mRNA molar fractions (*χ*^*mR*^ in mol mol_mR_^−1^) and mass fractions (*ϕ*^*mR*^ in g g_mR_^−1^) by accounting for transcript length before COG-level aggregation. Details of the biomass reconstruction and omics processing are provided in *Methods* and the *Supplementary Information*.

Across compiled physiological datasets, the RNA-to-protein mass ratio 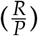 increases consistently with the specific growth rate *µ* (Figure 2.1a). This relationship provides a useful physiological coordinate because it links faster growth to increased allocation toward RNA-rich, ribosome-associated biomass [39, 26, 10]. The Dai et al. data show two approximately linear branches, consistent with earlier interpretations that slow-growth cells can maintain ribosome excess or storage-like ribosome capacity under nutrient-limited conditions [26, 10]. In this study, the 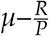 relationship was used to estimate growth-rate-dependent RNA species, ribosome abundance, and translation-associated components for model initialization.

**Figure 2.1.**
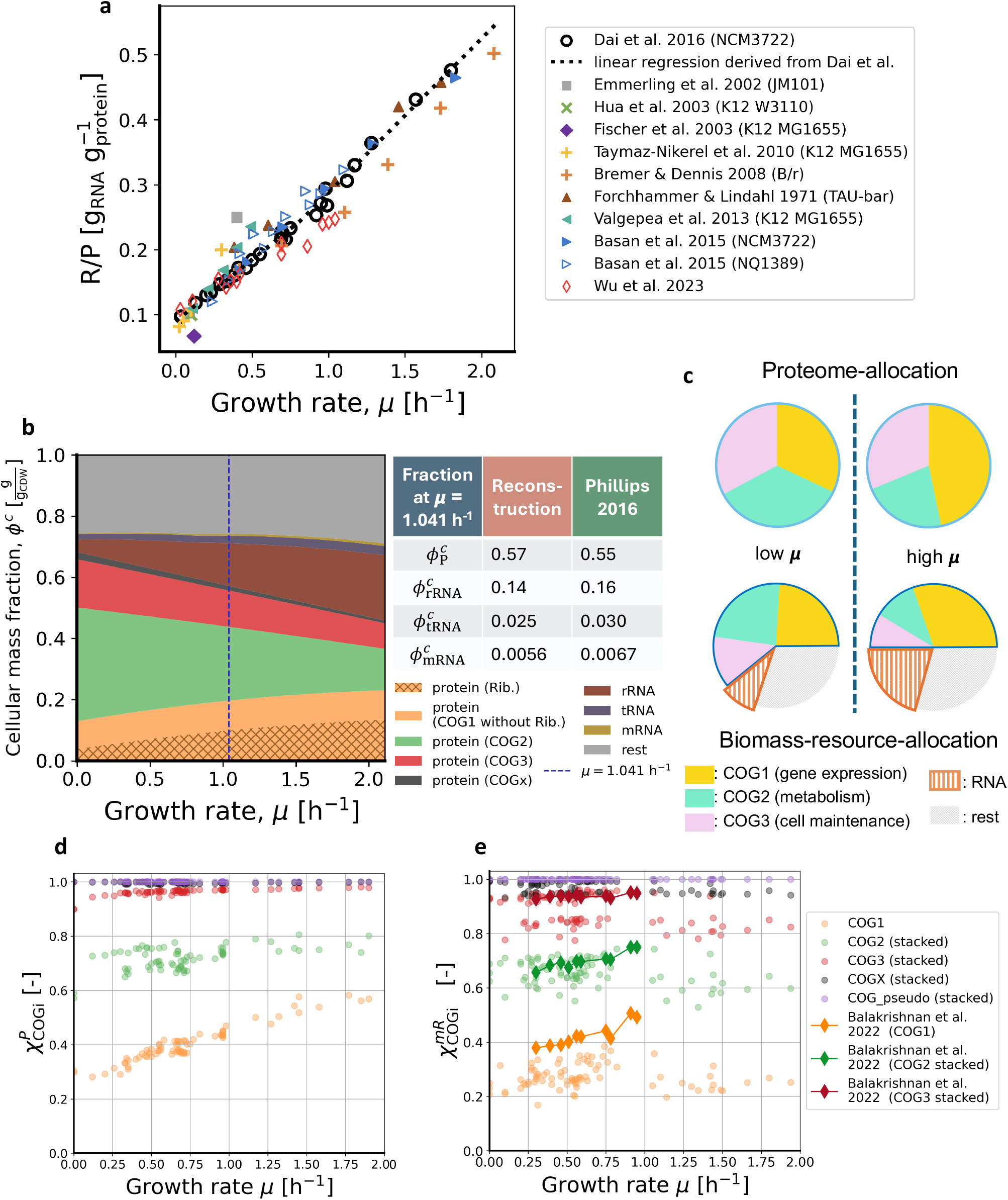
Growth-rate-dependent physiological and omics landscape of cellular resource allocation in *Escherichia coli*. **(a)** RNA-to-protein mass ratio 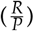 as a function of growth rate. Data points were collected from physiological studies of different *E. coli* strains and growth conditions. The dotted line shows the piecewise linear relationship fitted to the measurements of Dai et al. [12, 23, 14, 43, 6, 15, 46, 3, 10, 47]; **(b)** Reconstructed growth-rate-dependent biomass composition expressed as mass fractions relative to cell dry weight. The total protein fraction follows a growth-rate-dependent fit to compiled measurements [43, 12, 23, 14, 6] and is partitioned into COG1–3 and an unassigned COGx fraction using reprocessed proteomics datasets [38, 33, 48]. COG1 is further resolved into ribosomal proteins, shown by cross-hatching, and the remaining COG1 proteins. Total RNA is calculated from the fitted protein fraction and the Dai et al. 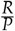 relationship [10]. mRNA is assigned a constant total-RNA mass fraction of 0.0329 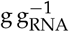 after calibration to the total-mRNA abundance scale reported by Balakrishnan et al. [2], and the remaining RNA is partitioned between tRNA and rRNA using a growth-rate-dependent tRNA-to-rRNA ratio [22]. The adjacent table compares the reconstructed fractions at *µ* = 1.041 h^−1^, indicated by the vertical blue dashed line, with reference biomass-composition values reportedby Milo and Phillips [30]; **(c)** Schematic comparison between proteome allocation and biomass-resource allocation; **(d)** COG-level proteome molar allocation derived from reprocessed proteomics datasets [38, 33, 48]; **(e)** COG-level transcriptome molar allocation derived from reprocessed transcriptomics datasets [13, 9, 37]. COG-level transcriptomic estimates from Balakrishnan et al. [2] are included for comparison.

**Figure 2.2.**
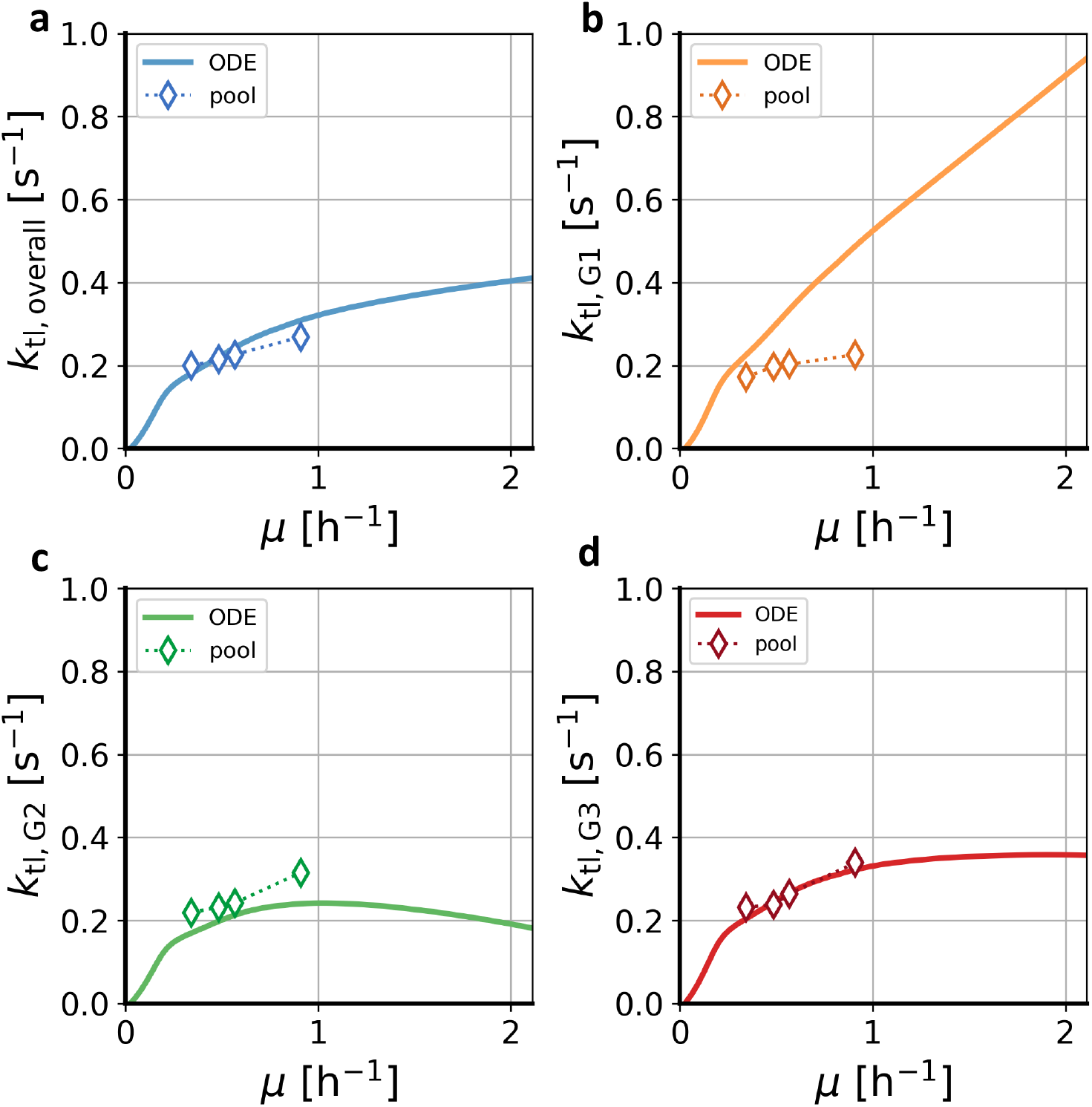
Growth-rate-dependent translation frequency of overall and COG-specific protein synthesis. **(a)** Simulated average translation frequency of the overall expressed protein pool as a function of growth rate. The solid line represents the ODE-based simulation, and diamond markers represent pool-based estimates derived from the matched proteomic and transcriptomic data of Balakrishnan et al. [2]; **(b–d)** COG-specific average translation frequencies for COG1, COG2, and COG3. COG1 represents gene-expression-related functions, COG2 represents metabolism-related functions, and COG3 represents cell-maintenance-related functions [38, 27]. Solid lines show ODE-based simulations, whereas diamond markers show pool-based estimates from the four selected datasets in Balakrishnan et al.’s work. The comparison provides an omics-derived reference for the growth-dependent translation-frequency trends obtained from the simulator.

The reconstructed biomass composition shows that the total protein fraction decreases mildly with growth rate, whereas the RNA fraction increases (Figure 2.1b). Over the quantified range from *µ* = 0.10 to 2.11 h^−1^, the protein mass fraction fitted by linear regression to compiled measurements (see *Methods*) decreases from approximately 0.672 to 0.458 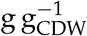, whereas 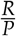 increases from approximately 0.109 to 0.551. The corresponding RNA mass fraction rises from approximately 0.073 to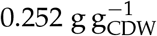. This increase is mainly attributable to rRNA, with tRNA and mRNA contributing much smaller fractions. Thus, the growth-rate-dependent rise in 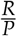 primarily reflects the expansion of ribosome-associated RNA rather than a uniform increase in all RNA species. Faster growth is therefore characterized by an RNA- and ribosome-enriched biomass state. The values reported by Milo and Phillips [30] were not used in the reconstruction and provide an external comparison at one growth rate. As shown in the table adjacent to Figure 2.1b, the reconstructed protein, mRNA, tRNA, and rRNA mass fractions are close to the corresponding reference values.

The biomass-resource-allocation view illustrated in the lower row of Figure 2.1c differs from the classical proteome-allocation view shown in the upper row. Proteome allocation describes redistribution within a constrained protein pool among functional sectors [39, 4, 33, 40]. In this framework, gene-expression-related proteins, including ribosomal proteins, and metabolism-related proteins are redistributed as the growth condition changes. Biomass-resource allocation instead uses total cell dry weight as the reference scale and captures simultaneous changes in protein, RNA, and the remaining biomass. It therefore accounts both for redistribution within the proteome and for the decline in the total protein fraction with increasing growth rate. This distinction is important because a ribosome-associated protein sector can increase strongly within the proteome while changing more moderately on the whole-biomass scale. Panel b makes this scale dependence explicit: the cellular mass fraction of the complete COG1 protein sector, which comprises gene-expression-related proteins, increases moderately with growth rate. Its ribosomal-protein subset, highlighted by cross-hatching, increases more strongly, whereas rRNA undergoes the largest expansion among the ribosome-associated biomass components.

The growth-rate-dependent molar fractions of the proteome and transcriptome were used in the subsequent translation-frequency analysis. Reprocessed proteomic datasets [38, 33, 48] show a clear redistribution within the protein pool (Figure 2.1d): the COG1 fraction increases with growth rate, the COG2 fraction decreases, and the COG3 fraction shows a weaker dependence. By contrast, the transcriptomic datasets [13, 9, 37] vary among studies and show no consistent monotonic growth-rate dependence (Figure 2.1e). Across these datasets, the mean mRNA molar fractions are 0.273 mol mol_mR_^−1^ for COG1, 0.382 mol mol_mR_^−1^ for COG2, and 0.229 mol mol_mR_^−1^ for COG3. The Balakrishnan et al. dataset [2] was included only for comparison and was not used to calculate the mean growth-dependent transcriptomic molar fractions used to initialize the ODE simulator. Relative to the other transcriptomic datasets, it shows a higher mean COG1 fraction (0.426 mol mol_mR_^−1^) and a lower mean COG2 fraction (0.276 mol mol_mR_^−1^) than the corresponding means of the other transcriptomic datasets. These between-study differences in transcript allocation must be considered when interpreting sector-specific translation estimates.

Together, these data define the growth-rate-dependent resource environment in which translation operates. Faster growth is associated with rRNA expansion, altered total protein allocation, redistribution of the proteome toward gene-expression-related proteins, and COG-specific changes in mRNA template availability. This integrated landscape provides the quantitative basis for the subsequent ODE-based analysis of mRNA–ribosome interaction, translation frequency, and ribosome utilization.

### 2.2 ODE-based simulation links resource allocation to translation behavior

The resource landscape described above specifies the growth-rate-dependent availability of ribosomes, RNA species, and COG-specific protein and mRNA pools. However, static abundance information does not determine how efficiently mRNA templates are translated. To connect resource allocation with translation dynamics, an ODE-based simulator was used to represent the major stages of bacterial translation, including initiation, elongation, termination, ribosome recycling, and tRNA charging. For each growth-rate condition, the simulator was initialized with the corresponding molecular abundances, represented as cellular volumetric component concentrations, whose units are written in µM or mM, which refers to µmol 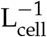 or mmol 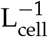 respectively, and solved to a steady-state distribution. Here, steady state means that component concentrations become approximately time-invariant while translation fluxes continue through the system.

The primary output was the average translation frequency, *k*_tl_, defined as the protein production rate normalized by the abundance of the relevant mRNA template pool:

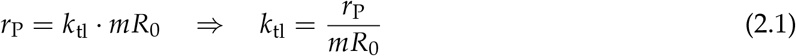

For COG-specific estimates, proteomic and transcriptomic molar fractions were used to assign protein production demand and mRNA template abundance to the three functional sectors.

Pool-based estimates were derived from four *E. coli* growth experiments reported by Balakrishnan et al. In each experiment, proteomic and transcriptomic measurements were obtained from the same cultivation sample, allowing the protein and mRNA pools to be compared for a matched strain and growth condition. The corresponding proteomic measurements are included in the dataset reported by Mori et al. [33], which also contributes to the growth-rate-dependent COG molar fractions in Figure 2.1d. The four measured growth rates are 0.343, 0.485, 0.568, and 0.91 h^−1^.

Under balanced exponential growth, the net protein production associated with biomass accumulation can be approximated as *µ* · *P* [13, 27, 2]:

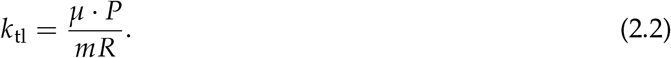

Here, *P* and *mR* denote protein and mRNA abundances on the same physiological scale. For the overall estimate, total protein and mRNA abundances were used; for each COG-specific estimate, the abundances of all proteins and mRNAs assigned to that COG group were summed separately before calculating their ratio. The pool-based estimates were used to assess whether the simulated *k*_tl_ reproduced a comparable magnitude and growth-dependent trend at the matched experimental conditions. This output-level comparison does not directly validate the individual kinetic constants in the ODE model, but it provides consistency between simulated results under a comprehensive kinetic setup and pool-based estimates from omics data.

The simulated overall translation frequency increases with growth rate in a sublinear manner (Figure 2.2a). Between *µ* = 0.10 and 2.11 h^−1^, *k*_tl_ increases from 0.046 to 0.410 s^−1^, an increase of approximately 784.8%. The curve rises steeply at low growth rates and more gradually at higher growth rates, indicating that the average protein production rate per mRNA template increases during faster growth but with diminishing incremental gains. At the four Balakrishnan reference conditions, the largest absolute simulation–reference difference is 0.040 s^−1^ at *µ* = 0.91 h^−1^ (14.8% of the reference value).

The COG-specific trajectories reveal that translation frequency is not uniform across functional sectors (Figure 2.2b–d). COG1 shows the strongest growth-dependent increase, from 0.051 to 0.940 s^−1^ over the same range. COG2 increases from 0.047 s^−1^ to a maximum of 0.242 s^−1^ at *µ* = 1.02 h^−1^ and then declines to 0.182 s^−1^. COG3 increases from 0.054 s^−1^ and approaches a plateau, reaching a maximum of 0.358 s^−1^ at *µ* = 1.90 h^−1^. The largest discrepancy between simulation and pool-based estimates occurs for COG1 (0.262 s^−1^ at *µ* = 0.91 h^−1^). Because the Balakrishnan dataset contains a relatively high molar fraction of COG1 transcripts (Figure 2.1e), a given COG1 protein-production demand near this growth rate can be supported by a lower average translation frequency per COG1 mRNA. This discrepancy is therefore best interpreted as a dataset-specific transcriptome feature rather than as a direct failure of the simulated trend.

### 2.3 mRNA–ribosome distributions are consistent with a limitation-regime crossover

The steady-state distributions of mRNA and ribosomes indicate how the sublinear increase in translation frequency emerges from molecular allocation (Figure 2.3). Total mRNA concentration increases with growth rate and follows the overall trend of the calibrated or re-estimated values from Valgepea et al. and Balakrishnan et al. [46, 2]. In the coarse-grained model, each mRNA unit is assigned one effective ribosome-binding site (RBS), such that the total concentration of modeled RBSs represents the total mRNA concentration. This pool is partitioned between occupied RBSs represented in mRNA-containing ribosomal complexes and free RBSs available for ribosome binding. The resulting binary partition describes initiation-site occupancy rather than the number of ribosomes distributed along the coding region of each transcript. The concentration of occupied RBSs remains relatively stable across most of the growth-rate range, whereas the free-RBS concentration increases strongly. Consequently, the fraction of occupied RBSs decreases as the total mRNA pool expands.

**Figure 2_3.**
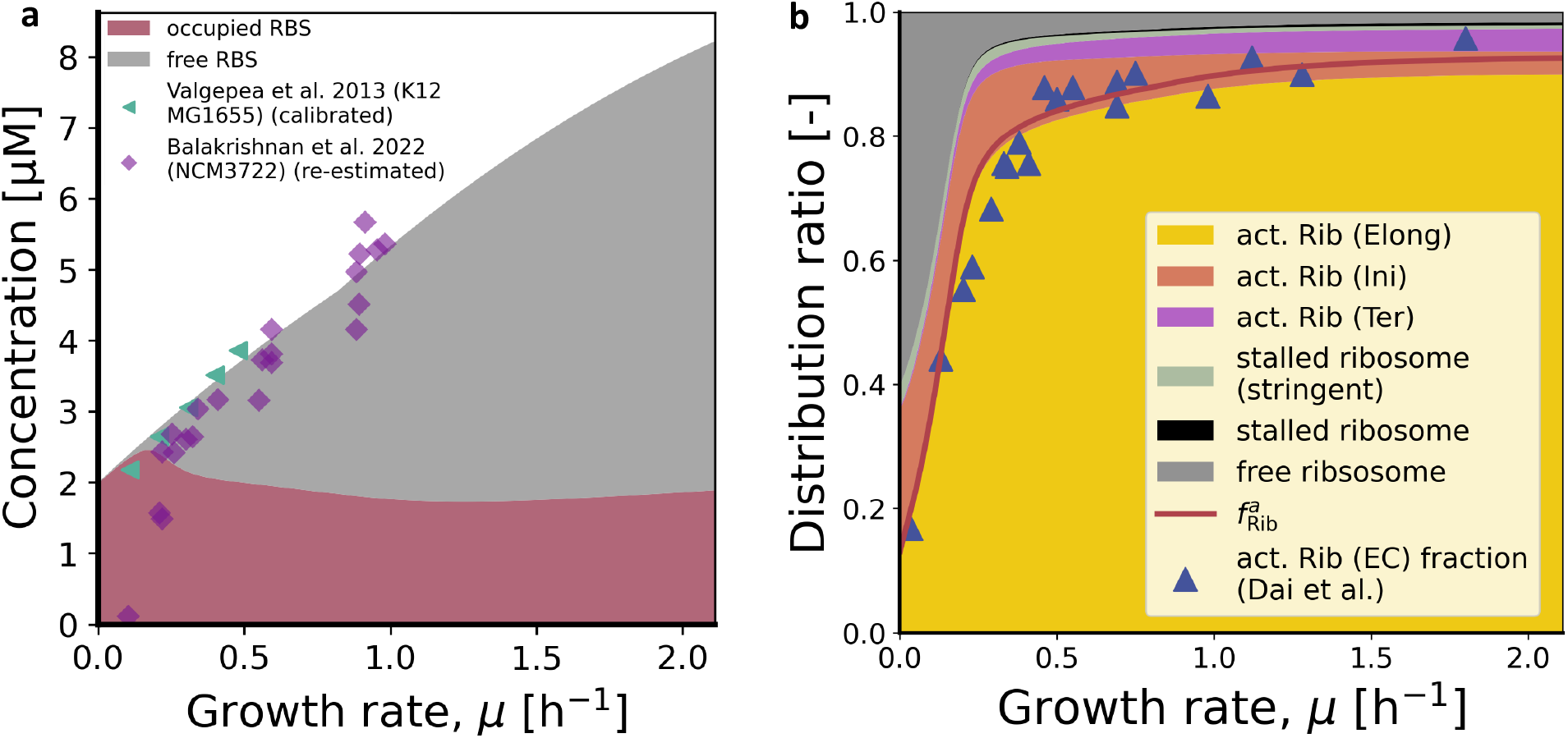
Steady-state distributions of mRNA and ribosome states. **(a)** Steady-state concentrations of occupied and free ribosome-binding sites (RBSs) as functions of growth rate. Each modeled mRNA unit contributes one effective RBS; therefore, the upper boundary of the stacked areas represents the total cellular mRNA concentration. Calibrated mRNA-abundance estimates from Valgepea et al. [46] and values re-estimated from Balakrishnan et al. [2] are shown for comparison. **(b)** Simulated steady-state ribosome distribution, partitioned among elongation, initiation, termination, stalled, and non-active states. The red curve shows the amino-acid- or nascent-chain-carrying ribosome fraction, 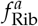, whereas the upper boundary of the elongation area represents the EC fraction. Blue triangles show the active-ribosome equivalents derived by Dai et al. [10].

Ribosome allocation follows a contrasting pattern. Between *µ* = 0.10 and 2.11 h^−1^, the fraction of ribosomes associated with initiation, elongation, and termination increases sharply before gradually approaching a high-growth plateau (Figure 2.3b). However, not all ribosomal complexes in these three phases carry an amino acid or nascent peptide. Early initiation states represent ribosome docking and initiation-complex assembly, whereas post-release termination states are involved in ribosome recycling. After excluding these non-amino-acid-carrying states, the remaining amino-acid-carrying active-ribosome fraction, 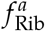, increases from 0.357 to 0.926. Over the same growth-rate range, the bound-RBS fraction, 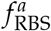, decreases from 0.979 to 0.230 (Figure 2.4a). The phase-resolved distribution further shows that most translation-associated ribosomes are present in elongation complexes (ECs), whereas initiation and termination complexes account for smaller fractions (Figure 2.3b). Stalled ribosome states remain minor components of the total ribosome pool.

**Figure 2.4.**
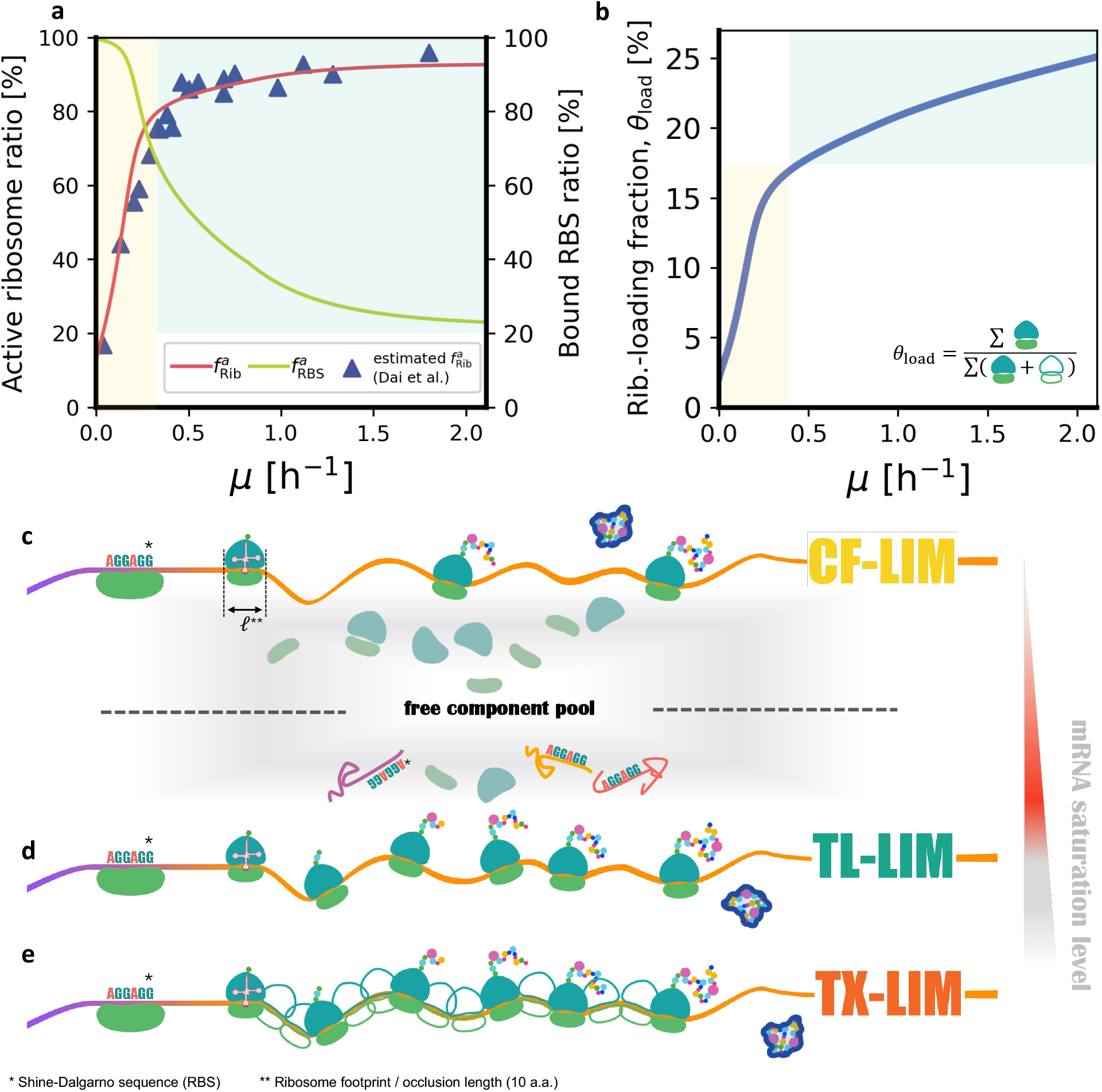
Molecular occupancy states and translation-limitation regimes. **(a)** Growth-dependent changes in active-ribosome and bound-RBS fractions. These allocation trends are consistent with a crossover from low-growth complex-formation limitation (yellow area) toward high-growth translation limitation (blue area). **(b)** Estimated mRNA ribosome-loading fraction, *θ*_load_ = *n*/*n*_max_, expressed as the percentage of the theoretical footprint-limited loading capacity that is occupied. This transcript-wide measure is distinct from RBS occupancy; the yellow and blue areas correspond to the same limitation regimes as in panel a. **(c–e)** Conceptual limiting cases adapted from Calabrese et al. [8]. **(c)** complex-formation limitation (CF-LIM), in which mRNA and ribosome availability jointly affect production and a substantial free-ribosome pool remains; **(d)** translation limitation (TL-LIM), in which abundant transcripts compete for a limiting ribosome pool and many RBSs remain free; and **(e)** transcription limitation (TX-LIM), in which near-saturated mRNA loading makes transcript abundance the principal limitation. TX-LIM is not reached over the simulated physiological range. The adjacent gradient indicates the mRNA-saturation level represented by the loading fraction in panel b. The gray interval between TL-LIM and TX-LIM denotes higher mRNA loading that is not reached in the present simulation. ^∗^ Canonical Shine–Dalgarno sequence representing the RBS. ^∗∗^ The ribosome footprint is the minimum coding-region spacing between adjacent mRNA-bound ribosomes imposed by ribosome occlusion.

The simulated 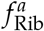 was compared with the active-ribosome equivalents reported by Dai et al. [10]. Dai et al. inferred an elongation velocity from the waiting time to the first detectable LacZ protein signal after correcting for the initiation delay. In the present model, elongation velocity refers specifically to repeated amino-acid incorporation within the EC cycle. If the experimentally inferred velocity represented only this process, the active-ribosome equivalent derived by steady-state mass balance would correspond most directly to the EC fraction. However, the completion-dependent LacZ signal does not resolve elongation from the terminal steps required before peptide release. The inferred active-ribosome equivalent therefore need not correspond exclusively to ribosomes in ECs. We consequently compared the Dai estimates with 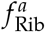, which includes amino-acid- or nascent-chain-carrying ribosomes in late initiation, elongation, and pre-release termination states. The simulated EC fraction and 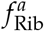 nevertheless remain close across the investigated growth-rate range (Figure 2.3b). The largest absolute difference between 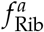 and the Dai reference values is 0.126 at *µ* = 0.23 h^−1^. Because the Dai elongation-velocity and active-ribosome estimates contributed to calibrating selected effective kinetic terms, this comparison serves as a calibration-consistency check.

RBS occupancy and transcript-wide ribosome loading describe different levels of organization. The bound-RBS fraction reports the fraction of mRNA initiation sites represented in mRNA-containing ribosomal complexes. By contrast, the mRNA ribosome-loading fraction, *θ*_load_, estimates the mean number of ribosomes distributed along a translated coding region relative to its theoretical footprint-limited maximum (see Figure 2.4 and *Methods*). An RBS can therefore be counted as occupied even when the remainder of the transcript carries substantially fewer ribosomes than the theoretical maximum. Details of the estimation approach are provided in *Methods*.

To interpret these allocation patterns, we use the limiting-regime framework proposed by Calabrese et al. [8]. In the complex-formation-limited regime (CF-LIM, Figure 2.4c), protein production depends jointly on mRNA availability, ribosome availability, and the formation of productive mRNA–ribosome complexes. In the translation-limited regime (TL-LIM, Figure 2.4d), transcripts are sufficiently abundant that ribosome availability becomes the principal limitation. In the transcription-limited regime (TX-LIM, Figure 2.4e), mRNA templates approach saturation with ribosomes, namely, the maximal *θ*_load_, and protein production depends primarily on transcript abundance. These regimes are used here as limiting-case interpretations of the simulated allocation and kinetic relationships, not as additional discrete states imposed on the ODE system.

Viewed within this framework, the mRNA and ribosome distributions are consistent with a growth-dependent crossover in translation limitation (Figure 2.4a–b). At low growth rates, scarce free RBSs, high RBS occupancy, and a large non-active ribosome pool are consistent with CF-LIM (yellow-shadowed area in Figure 2.4a), in which productive mRNA–ribosome complex formation remains an important determinant of output. As the growth rate increases, the free RBS pool expands, the non-active ribosome pool is depleted, and ribosomes are increasingly recruited into elongation. This allocation pattern is consistent with an approach toward TL-LIM (blue-shadowed area in Figure 2.4a), in which ribosome availability and utilization dominate once mRNA-binding capacity is abundant. These distributions motivate the regime interpretation; the immediately following production-rate analysis tests it against the simulated dependencies on free RBS and total ribosome concentrations.

The transcript-wide mRNA ribosome-loading fraction provides a separate test of whether the simulated system approaches TX-LIM. The model-derived *θ*_load_ increases from 0.067 (6.7%) at *µ* = 0.10 h^−1^ to 0.251 (25.1%) at *µ* = 2.11 h^−1^, but remains far below the theoretical maximum of one (Figure 2.4b; Equation 4.20 in *Methods*). The simulation, therefore, does not support a near-saturated, transcription-limited state over the physiological growth-rate range. This loading fraction is a coarse-grained model-derived indicator rather than a direct measurement of polysome occupancy. Within the model, higher translation output is achieved while substantial transcript-wide loading capacity remains unused, rather than requiring near-maximal ribosome packing on mRNA.

### 2.4 Production kinetics support a continuous limitation-regime crossover

The overall protein production rate provides the kinetic evidence for the limitation-regime interpretation introduced above. In a model-specific adaptation of the CF-LIM and TL-LIM limiting cases, protein production can be summarized by a Michaelis–Menten-like dependence on available mRNA-binding capacity and an approximately proportional dependence on total ribosome abundance:

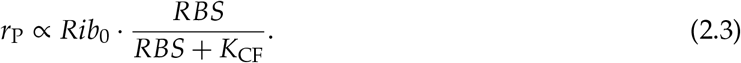

Here, *Rib*_0_ denotes total ribosome concentration, *RBS* denotes free ribosome-binding-site concentration, and *K*_CF_ is an effective half-saturation constant for mRNA–ribosome complex formation. When free RBSs are scarce, the unsaturated RBS-dependent term varies strongly with *RBS*; protein production therefore depends on both available transcript sites and ribosome abundance, as expected for CF-LIM. As free-RBS availability increases, this term approaches saturation and *r*_P_ becomes increasingly proportional to *Rib*_0_, which is the limiting behavior expected when the system approaches TL-LIM. The two regions in Figure 2.5a therefore represent low- and high-*RBS* branches of the same continuous growth-dependent trajectory rather than two separately simulated kinetic models.

**Figure 2.5.**
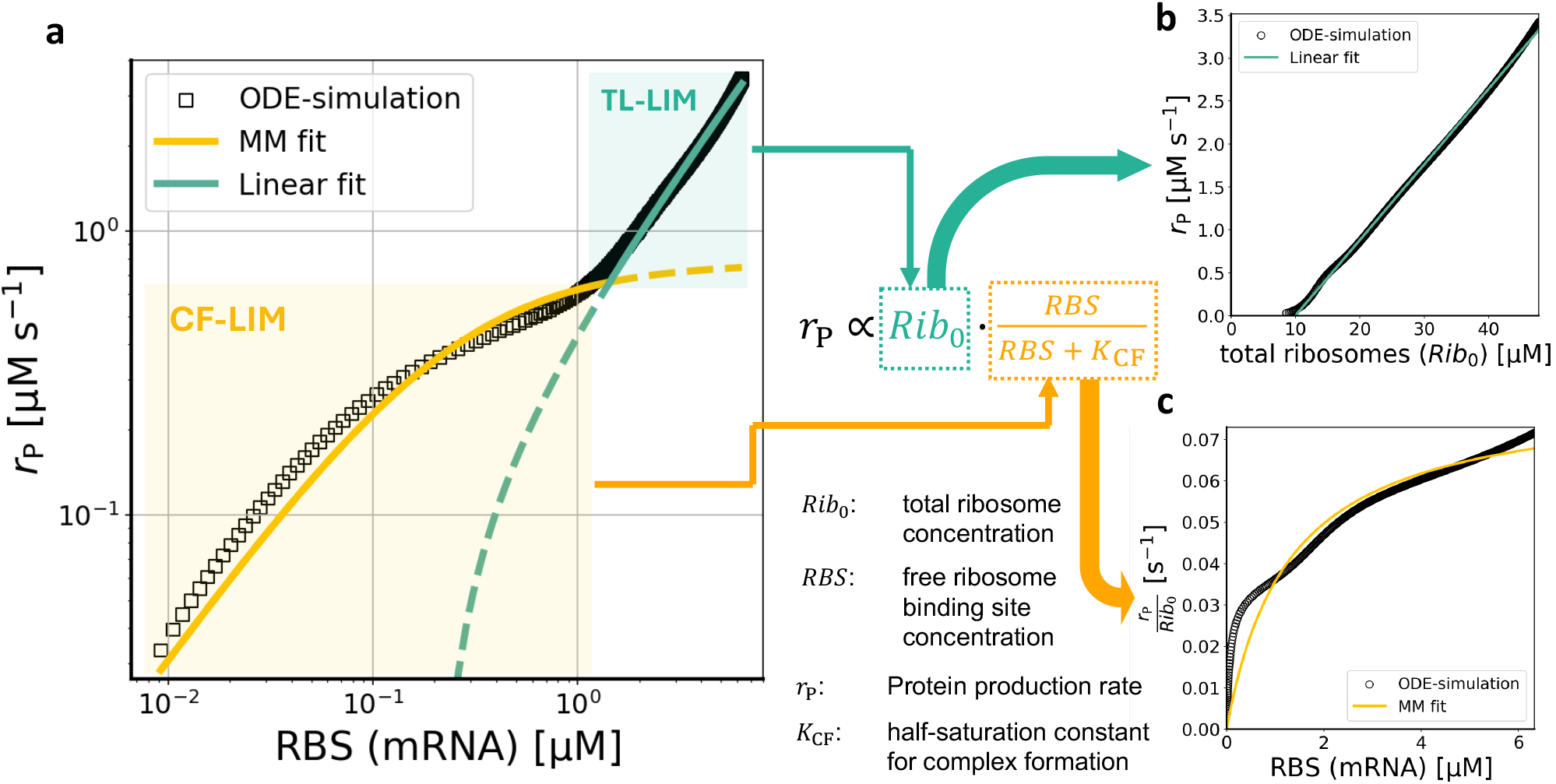
Production-rate relationships supporting a continuous limitation-regime crossover. **(a)** Protein production rate (*r*_P_) as a function of free ribosome-binding-site concentration (*RBS*), used here as a simulator-specific measure of available mRNA-binding capacity. The low-*RBS* branch was summarized by a Michaelis–Menten (MM) fit and the high-*RBS* branch by a linear fit; the split at *RBS* ≃ 1.442 µM minimized their combined residual sum of squares. The shaded regions denote CF-LIM-consistent and TL-LIM-consistent branches of one continuous simulated trajectory, not discrete states imposed on the ODE system; **(b)** Approximately linear relationship between *r*_P_ and total ribosome concentration *Rib*_0_, supporting an approximately proportional dependence of the production scale on ribosome abundance; **(c)** Saturating relationship between ribosome-normalized protein production rate 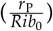 and free RBS concentration, showing diminishing production gains per ribosome as free-RBS availability increases. Panels b and c provide the direct model-internal kinetic basis for interpreting the high-*RBS* branch in panel a as an approach toward TL-LIM.

To describe the change in curvature, the low-*RBS* branch was fitted with an MM function and the high-*RBS* branch with a linear function. The residual-minimizing split occurred at *RBS* ≃ 1.442 µM, with *R*^2^ = 0.9729 for the MM branch and *R*^2^ = 0.9945 for the linear branch. The apparent linear dependence on *RBS* in the high branch does not imply that RBS availability remains the principal limitation, because free RBS and total ribosome concentrations increase together along the simulated growth trajectory.

The TL-LIM interpretation is supported by the approximately linear relationship between *r*_P_ and *Rib*_0_ (*R*^2^ = 0.9988; Figure 2.5b). The fitted line has an extrapolated *Rib*_0_-axis intercept of approximately 10 µM, indicating that protein production approaches zero while a residual ribosome pool remains in the model. This behavior is consistent with reports that slow-growing cells maintain excess or storage-like ribosome capacity [26, 10], although the fitted intercept is not a direct measurement of such a pool. The complementary relationship between 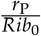 and *RBS* is saturating (*R*^2^ = 0.9381; Figure 2.5c), showing that additional free RBSs provide diminishing production gains per ribosome, whereas total ribosome abundance continues to determine the production scale. The free-RBS concentration is a simulator-specific measure of available transcript binding capacity and is not identical to the total mRNA concentration in the original Calabrese formulation [8]. Together, these fits provide model-internal evidence for a continuous crossover from CF-LIM toward TL-LIM, rather than an independent experimental classification or a discrete regime boundary.

### 2.5 Initiation kinetics and factor availability

The preceding analysis relates protein production to ribosome abundance and free-RBS availability at the system level, thereby connecting the simulated trajectory to the CF-LIM and TL-LIM framework. These macroscopic relationships describe how overall molecular allocation and protein production covary, but they do not identify the elementary kinetic steps through which the translation flux is established. We therefore resolved the simulated translation frequency at the productive-initiation boundary before examining the contributions of individual initiation-side components.

The translation frequency *k*_tl_ is calculated from the net retained protein production rate and therefore includes the opposing contribution of protein degradation. It is not directly comparable to the frequency of initiation events, which contribute to nascent protein synthesis before degradation. To isolate the synthesis process, the modeled degradation flux was added back to define the nascent translation frequency

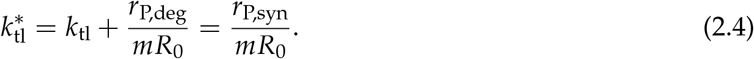

Here, *r*_P,deg_ is the modeled protein degradation flux and *r*_P,syn_ is the nascent protein synthesis flux. In Kremling’s kinetic formulation, productive initiation frequency is theoretically equivalent to 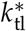 under steady-state translation because each successfully initiated translation cycle contributes one nascent protein product [28, 27]. The present simulator explicitly represents the sequential initiation intermediates, allowing this equivalence to be tested from independently calculated outputs.

As illustrated by the initiation pathway in Figure 2.6a, 70*SPIC* is the final initiation intermediate immediately preceding productive elongation, and *k*_i2_ is its first-order forward clearance constant. Their product therefore gives the flux completing initiation, *r*_ini_ = *k*_i2_ · 70*SPIC*. Normalization by the initial mRNA abundance gives the initiation frequency

**Figure 2.6.**
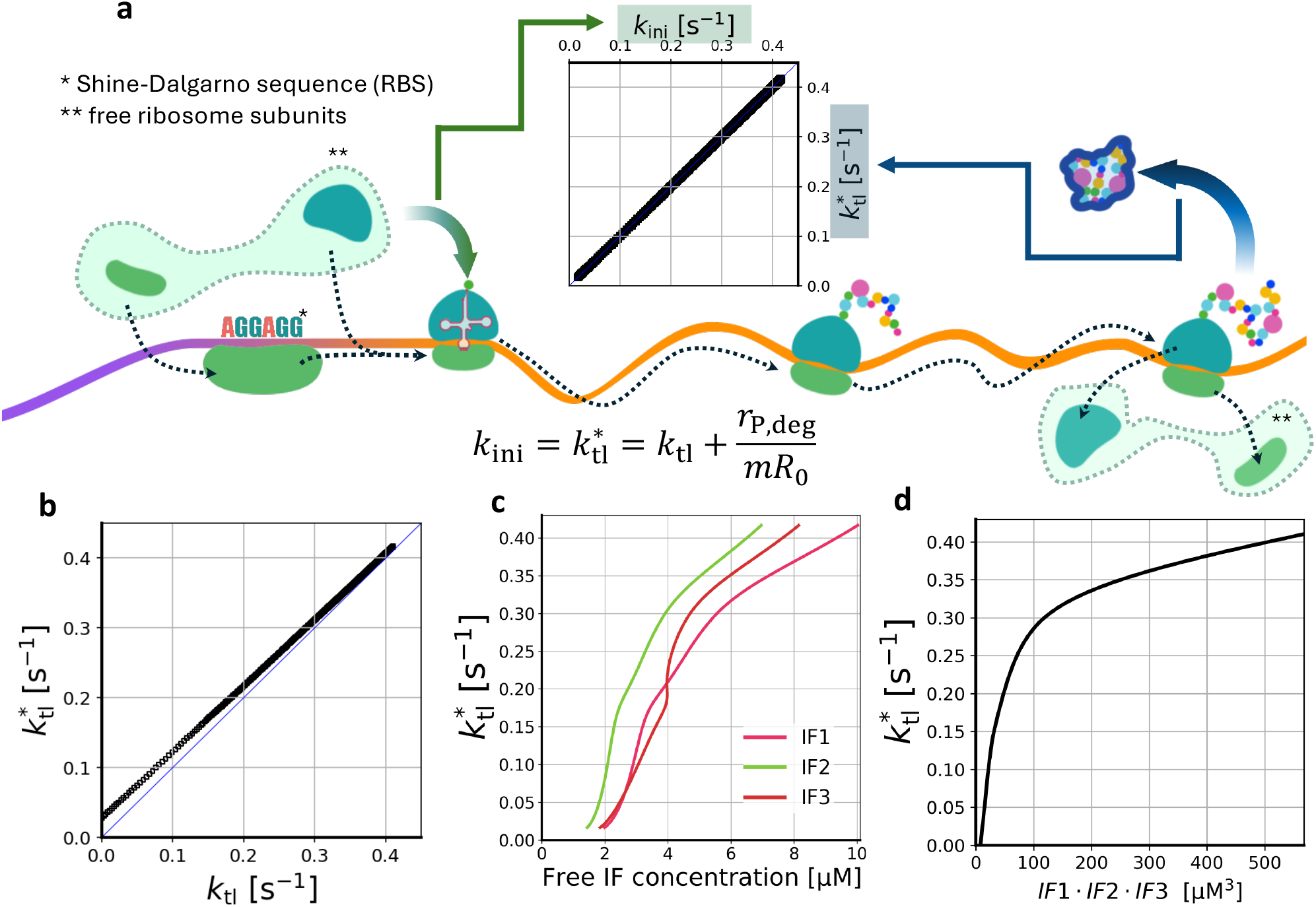
Initiation-side consistency and factor-availability relationships. **(a)** Schematic representation of translation initiation and comparison of initiation frequency with nascent translation frequency. Clearance of the final initiation intermediate, 70*SPIC*, with the forward rate constant *k*_i2_ gives *r*_ini_ = *k*_i2_ · 70*SPIC* and *k*_ini_ = *r*_ini_/*mR*_0_. The nascent translation frequency is defined as 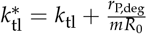 and is expected to equal *k*_ini_ at steady state. **(b)** Comparison between *k*_tl_ and 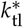; the blue identity line indicates equality between the two quantities. **(c)** Relationship between 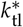 and the free concentrations of IF1, IF2, and IF3. **(d)** Relationship between 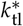 and the product of the free initiation-factor concentrations.

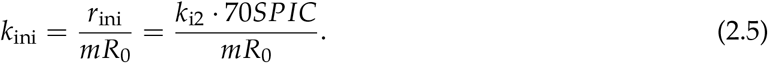

The simulated *k*_ini_ aligns closely with 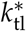 across growth rates (Figure 2.6a). Across the complete simulated grid, their maximum absolute difference is 9.1 *×* 10^−6^ s^−1^. This agreement verifies the expected steady-state flux equivalence and shows that nascent translation throughput can be represented by the productive initiation flux in the implemented model. It therefore provides the basis for examining which initiation-side states and factors accompany changes in translation frequency. However, flux equivalence alone does not establish initiation as the sole rate-limiting stage, because downstream translation steps can also affect throughput through ribosome occupancy and recycling.

The difference between *k*_tl_ and 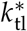 is most visible at low growth rates (Figure 2.6b). The corrected frequency increases from 0.071 s^−1^ at *µ* = 0.10 h^−1^ to 0.417 s^−1^ at *µ* = 2.11 h^−1^. The degradation correction shifts the trajectory above the identity line 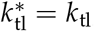, showing that protein degradation has a stronger relative effect under slow-growth conditions.

Having established *k*_ini_ as an equivalent steady-state readout of 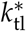, initiation-factor availability is next examined as a model-internal correlate of translation output. In the model, coordinated IF-dependent progression is represented by a coarse-grained mass-action term proportional to the product of free IF1, IF2, and IF3 concentrations:

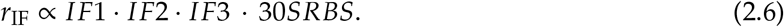

This expression should not be interpreted as an experimentally resolved elementary reaction in which all three factors bind simultaneously. Instead, it is an effective availability term summarizing the requirement that IF1, IF2, and IF3 jointly support efficient initiation-complex assembly and fidelity [1, 19].

The corrected nascent translation frequency increases with the free concentrations of IF1, IF2, and IF3 (Figure 2.6c), and also with the combined concentration product *IF*1 · *IF*2 · *IF*3 (Figure 2.6d). The relationship is steep at low IF availability and progressively flatter at higher values, indicating diminishing returns. Because the effective kinetic term in Equation 2.6 already encodes IF-dependent progression, these trends document how that implemented dependence behaves across the simulated range; they do not independently establish a causal role for IF availability.

The steady-state distributions of initiation intermediates and initiation factors indicate how this kinetic behavior emerges inside the simulator (Figure 2.7). At low growth rates, the early 30S–RBS complex is the dominant initiation intermediate, indicating that ribosomes can bind mRNA but progression toward downstream productive initiation complexes is limited. With increasing growth rate, occupancy shifts toward later initiation intermediates, suggesting more efficient progression through the initiation pathway.

**Figure 2.7.**
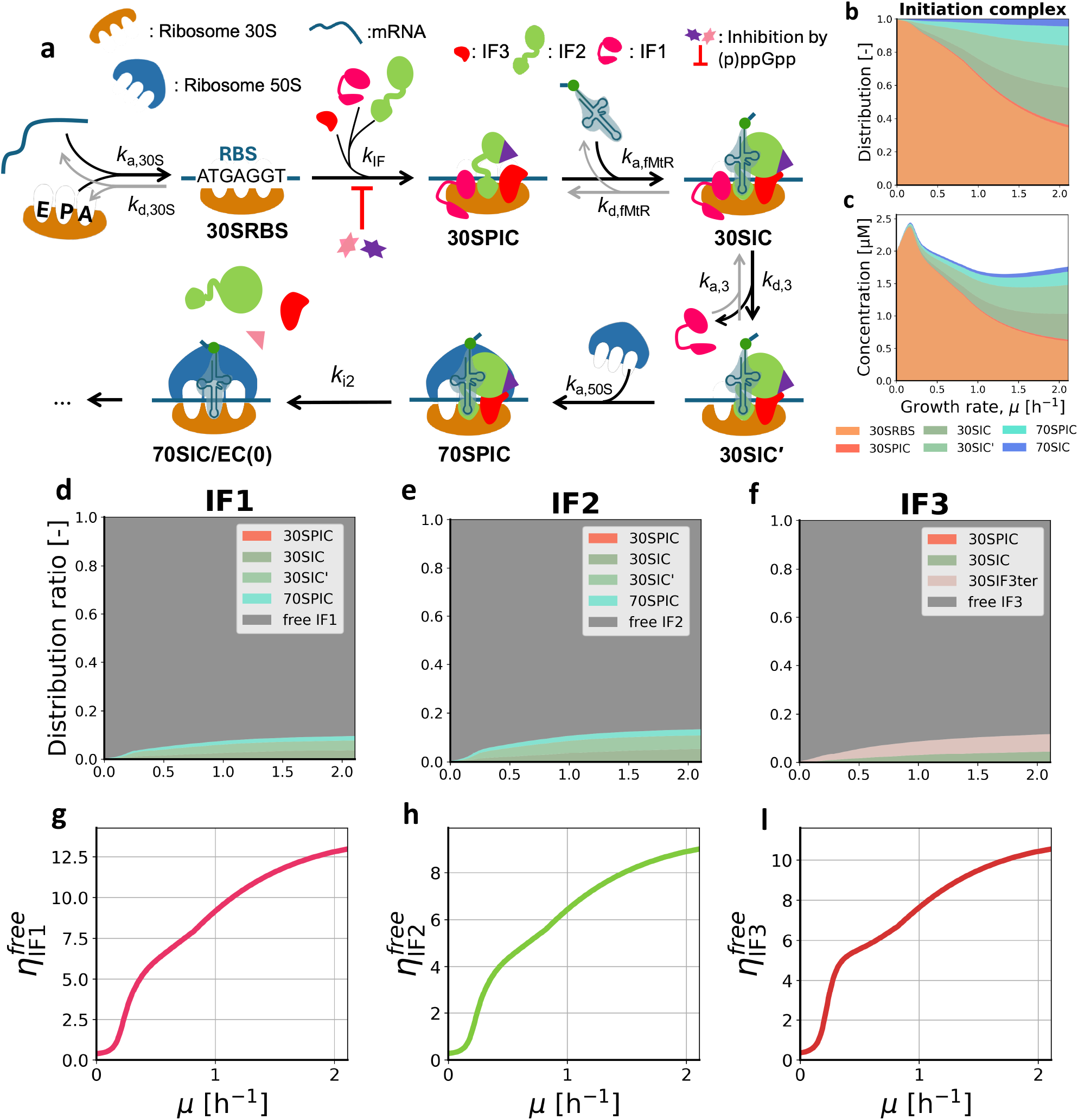
Steady-state distributions of initiation complexes and initiation factors across growth rates. **(a)** Schematic representation of the translation-initiation pathway implemented in the ODE simulator; **(b–c)** Steady-state distribution of initiation intermediates as a function of growth rate; **(d–f)** Allocation of IF1, IF2, and IF3 among free and complex-bound states; **(g–i)** Stoichiometric ratios of free IF1, IF2, and IF3 to free ribosomes. The increasing ratios, 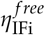, indicate that, despite depletion of free ribosomes at higher growth rates, a relative surplus of free initiation factors is maintained.

Although the amounts of IF1, IF2, and IF3 bound in initiation complexes increase with growth rate, the bound fractions remain small relative to the total pools. Thus, most initiation-factor molecules remain free at steady state. Between *µ* = 0.10 and 2.11 h^−1^, the free-IF-to-free-ribosome ratios, 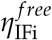, increase from 0.57 to 12.96 for IF1, from 0.41 to 9.00 for IF2, and from 0.57 to 10.54 for IF3. This behavior differs from ribosomes, which become increasingly recruited into active translation as growth rate increases. The increasing ratios show that the IF pools do not become depleted in parallel with the free-ribosome pool. However, this relative IF surplus does not by itself establish that IF availability causes the observed increase in initiation flux. Together with the implemented IF-dependent kinetic term, the preserved availability identifies a model-internal route by which effective initiation flux can be maintained as free ribosomes become scarce.

Together, the kinetic relationships and steady-state component distributions indicate that growth-dependent translation frequency is consistent with control by ribosome and mRNA availability, together with the modeled capacity of initiation factors to convert available mRNA–ribosome encounters into productive initiation events.

### 3 Discussion

This study quantitatively connects bacterial growth-rate-dependent resource allocation to mechanistic translation dynamics. The central result is that translation frequency is not determined solely by ribosome abundance. Instead, it emerges from the coupled availability of mRNA templates, ribosome recruitment and utilization, and the modeled IF-dependent progression of early mRNA–ribosome interactions. This interpretation extends classical growth-law theory from a primarily proteome-sector perspective toward a biomass- and kinetics-aware view of translation control.

The biomass-level reconstruction highlights an important distinction from standard proteome allocation. Classical growth-law theory describes how limited protein resources are redistributed among growth-rate-dependent sectors, with ribosome-associated proteins increasing as growth accelerates and metabolic sectors compensating accordingly [39, 26, 33, 40]. From a biomass-resource-allocation perspective, expressing total protein, rRNA, tRNA, mRNA, and remaining biomass on a common cell-dry-weight scale shows that faster growth is also characterized by a pronounced expansion of RNA, especially rRNA. This distinction matters because COG1 comprises proteins assigned to gene-expression-related functions, including ribosomal proteins, whereas ribosome biomass also contains a substantial rRNA contribution. Consequently, the growth-dependent increase in the COG1 fraction within the proteome is moderated on the whole-biomass scale by the concurrent decline in the total protein mass fraction. Conversely, the expansion of rRNA provides a biomass-level indicator of increased allocation to ribosomes. Proteome allocation and biomass-resource allocation are therefore complementary: the former resolves redistribution within the protein pool, whereas the latter captures the broader shift among protein, RNA, and other cellular biomass components.

The ODE simulation adds the dynamic layer that static resource allocation cannot provide. The simulated translation frequency increases sublinearly with growth rate, accompanied by higher ribosome utilization and decreasing free-ribosome availability. This behavior is consistent with the active-ribosome reference from Dai et al. used during calibration [10]. The comparison with Dai et al.’s data is therefore interpreted as a calibration-consistency check rather than as independent validation.

The COG-specific results further suggest that translation frequency differs among functional sectors. The strongest simulated increase is observed for COG1, whereas COG2 and COG3 show weaker or more saturating patterns. This provides a dynamic interpretation of omics allocation: functional sectors differ not only in protein and mRNA abundance, but also in the average translation frequency required to maintain sector-specific protein output. The comparison with Balakrishnan et al. is informative because that dataset contains four matched proteomic and transcriptomic measurements from the same *E. coli* growth experiment [2]. However, the lower pool-based estimate of *k*_tl,G1_ is strongly influenced by the relatively high molar fraction of COG1 transcripts in that dataset. Because Balakrishnan et al. used strain NCM3722, whereas several other transcriptomic datasets used strain MG1655 or related backgrounds [9, 13, 37], the COG1 discrepancy may partly reflect strain-specific transcriptome structure. Genetic and physiological differences between NCM3722 and MG1655 have been reported previously [7, 3]. Thus, the elevated COG1 transcript fraction should be treated as a dataset- and strain-specific feature rather than as a universal allocation pattern.

The limitation-regime framework proposed by Calabrese et al. provides a system-level interpretation of how mRNA-binding capacity and ribosome abundance shape the simulated protein-production trajectory [8]. Across the investigated growth range, the allocation and production-rate relationships are consistent with a continuous shift from CF-LIM-like toward TL-LIM-like behavior. As transcript-binding capacity expands, additional free RBSs yield diminishing production gains, whereas total ribosome abundance increasingly sets the production scale. To connect this system-level shift to initiation kinetics, we compared the degradation-corrected translation frequency 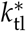 with the independently calculated initiation frequency *k*_ini_. Their close agreement links nascent protein synthesis to flux through the final initiation step, as expected from Kremling’s theoretical relationship [28, 27]. At the same time, the abundance of free IF1, IF2, and IF3 relative to free ribosomes increases as free ribosomes become scarce. The model summarizes the coordinated contribution of these factors through the effective concentration product *IF*1 · *IF*2 · *IF*3, consistent with evidence that all three factors contribute to the kinetics and fidelity of bacterial initiation [5, 1, 19, 17]. Rather than defining a separate limitation mechanism, this initiation-side behavior suggests how flux through the initiation pathway can be maintained as the system shifts from joint control by mRNA–ribosome binding toward stronger limitation by ribosome availability. The two analyses therefore provide complementary views: the regime-level analysis identifies how the principal resource constraint changes, whereas the initiation-side analysis describes how available mRNA–ribosome encounters are converted into translation flux. On this basis, we speculate that, as growth conditions become more favorable, *E. coli* may produce mRNA transcripts in excess of the immediate ribosome-binding demand. The associated decrease in fractional RBS occupancy could enable more complete utilization of the available ribosome pool. Under this hypothesis, producing additional mRNA would require less resource investment than assembling functional ribosomes, making excess transcript-binding capacity a potentially economical means of maximizing ribosome utilization. This interpretation is a biological hypothesis suggested by the simulated allocation pattern rather than a mechanism directly tested by the present model.

The central mechanistic limitation is the coarse-grained treatment of early mRNA engagement and IF-dependent progression. In the present model, 30S–mRNA/RBS engagement precedes a lumped IF-dependent transition and should be interpreted as an early 30S–mRNA engagement state rather than as a fully resolved productive 30S initiation complex. Structural and kinetic studies indicate that bacterial initiation is not governed by a single obligatory assembly order. mRNA and initiator tRNA can bind the 30S subunit through flexible routes, and initiation factors influence mRNA positioning as well as complex maturation [19]. A more detailed future model could represent parallel IF-independent and IF-assisted pathways, in which productive mRNA positioning depends on both free 30S subunits and IF-primed 30S complexes. Such a model would better reflect molecular initiation mechanisms but would require additional kinetic parameters for IF2/IF3 binding, IF1-dependent stabilization, mRNA relocation, initiator-tRNA recruitment, and 30S complex locking.

Several broader limitations define the current scope of the study. First, the physiological and omics datasets were compiled from different strains, media, platforms, and normalization procedures. This is particularly important for transcriptomic COG fractions and total protein mass fractions, both of which can vary across strain backgrounds and growth conditions. Second, COG classes are based on orthology and broad functional annotation rather than directly on growth-rate-dependent expression behavior [42]. COG1, COG2, and COG3 should therefore be interpreted as functional approximations of growth-law sectors, not as exact dynamic sectors. Future work could use expression-derived sector definitions, such as those proposed by Mori et al. [33], or introduce gene-specific translation parameters. Third, physiological scaling parameters such as cell dry weight-to-volume ratio and total protein fraction are unlikely to be invariant across all growth conditions, suggesting that more refined scaling will be required before a broadly generalizable translation-resource law can be established.

Despite these limitations, the model provides a useful quantitative bridge between growth-law resource allocation and translation-cycle kinetics. It suggests that the growth-dependent increase in translation frequency can be understood as an emergent systems property within the present simulator: mRNA availability reduces complex-formation limitation, ribosome recruitment increases translation capacity, and initiation-factor availability supports flux through the initiation pathway as free ribosomes become limiting. This framing can guide future extensions that incorporate elongation-factor availability, codon- or amino-acid-specific tRNA charging, and gene-specific translation dynamics.

## 4 Methods

### 4.1 ODE-based translation simulator

Before constructing the ODE input states, we reconstructed a growth-rate-dependent biomass-composition landscape for *Escherichia coli*. The reconstruction combined three classes of data sources. First, literature-derived physiological relationships were used to link growth rate, RNA-to-protein mass ratio 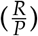, cell dry weight (CDW), and cellular volume [6, 3, 10, 12, 23, 14, 43, 15, 46, 47, 27]. A piecewise linear relationship fitted to the measurements of Dai et al. was used to map 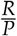 to *µ* over two growth-rate ranges (*Supplementary* Equation S13). This empirical 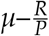 relationship defined the reference growth-rate coordinate of the simulator. In figures plotted against growth rate, each simulated condition was positioned at the reference *µ* corresponding to its prescribed 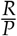 value. The growth-rate coordinate in these figures should therefore be distinguished from the independently calculated output *µ*_simulated_.

Second, biomass-level protein measurements were compiled to estimate the growth-dependent total protein mass fraction on a CDW scale, 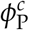, by linear regression [12, 23, 14, 43]. The total RNA mass fraction was then calculated as 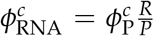 and partitioned into mRNA, rRNA, and tRNA. The residual CDW fraction, 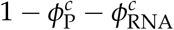, represents the remaining biomass components, including DNA, lipids, and peptidoglycan. The component-specific calculations are described below and extended in *Supplementary* section 1.4. Third, proteomic and transcriptomic datasets were reprocessed to assign translation-related protein pools and mRNA template pools to COG-level functional sectors [38, 33, 48, 13, 9, 37, 2].

All quantities were converted to common reference scales before use: global biomass quantities were expressed relative to CDW, and protein and transcriptomic allocations were converted into molar quantities where required for simulator initialization. This reconstruction defines the growth-rate-dependent abundances of ribosomes, mRNA templates, tRNAs, translation factors, enzymes, and small-molecule inputs used in the simulations.

Given these reconstructed resource constraints, an ODE-based simulator was used to represent the major kinetic steps of translation and to compute the steady-state distribution of molecular species. Because these components interact through a coupled nonlinear reaction network, the steady-state translation frequency (*k*_tl_) was obtained numerically from the ODE solutions rather than calculated from a single explicit algebraic expression.

The model should be interpreted as a quasi-steady-state resource-allocation simulator rather than as a literal time-course model of cellular growth. For each simulated condition, total cellular abundances were specified from empirical or fitted growth-dependent relationships, and the ODE system was integrated until the non-protein molecular species approached a steady distribution. The resulting state was then used to calculate translation frequency, initiation frequency, ribosome utilization, and related output quantities.

#### 4.1.1 Translation-cycle representation and ODE structure

The simulated translation system is organized into initiation, elongation, and termination modules, as summarized in Figure 4.1. Each module is represented by mass-action reactions or effective kinetic terms. The full simulator also includes tRNA charging, EF-Tu/EF-Ts-mediated nucleotide exchange, ribosome stalling, and static modifiers of selected GTP-dependent factor-association steps. These auxiliary modules close the translation cycle but are not analyzed as primary results because the analysis focuses on mRNA–ribosome interaction, ribosome utilization, and initiation-centered translation frequency. The complete ODE system is provided in *Supplementary* Table S1, and the extended initial-value calculations are described in *Supplementary* section 1.4.

**Figure 4.1.**
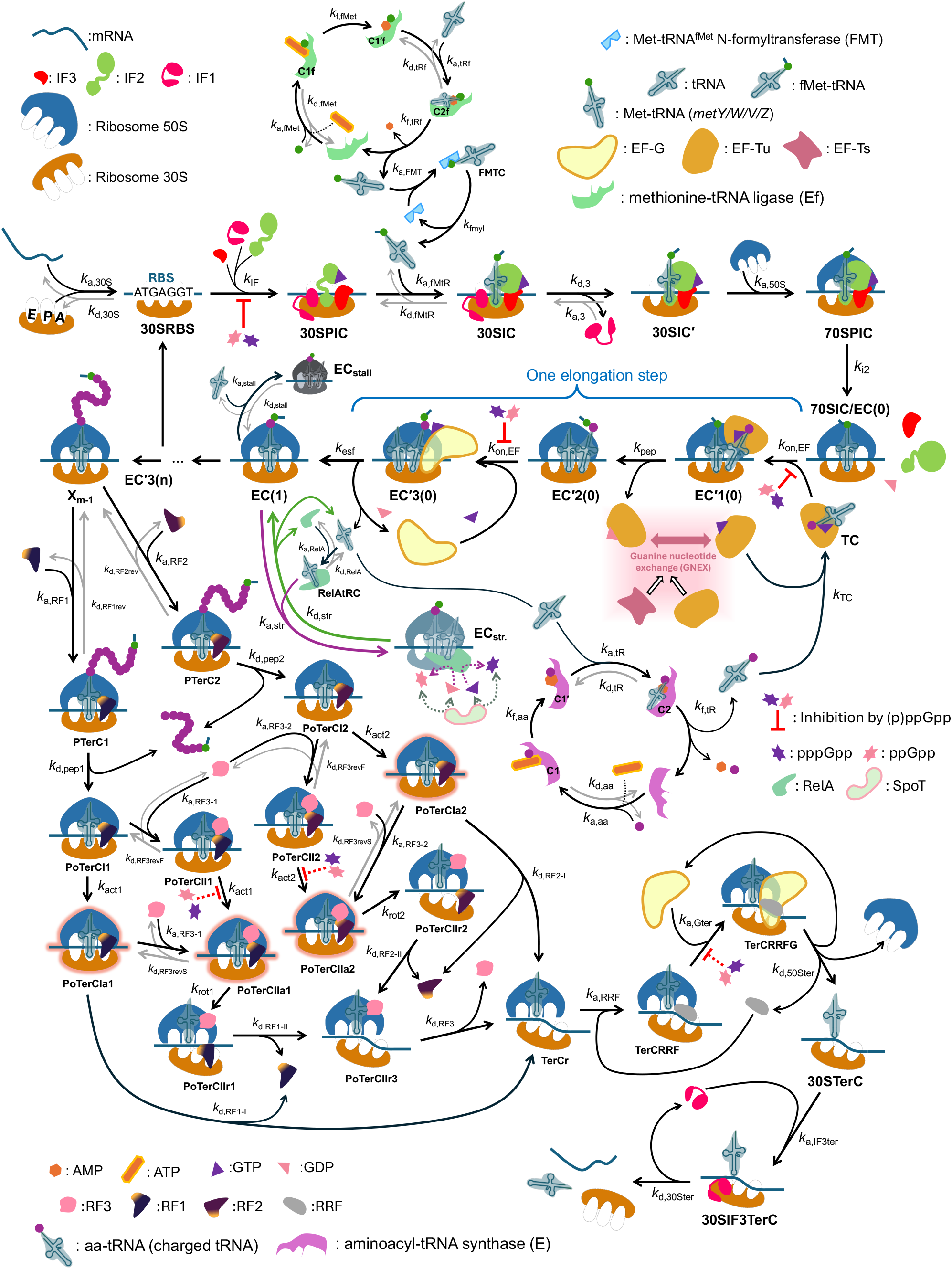
Translation-cycle reactions represented in the ODE simulator, including initiation, elongation, termination, tRNA charging, nucleotide exchange, and ribosome-stalling states.

##### Initiation stage

Translation initiation begins with binding of the 30S ribosomal subunit to an mRNA ribosome-binding site (RBS). In the model, the early 30S–RBS state is followed by an effective IF-dependent progression step involving IF1, IF2, and IF3, forming the 30S pre-initiation complex (30SPIC). This complex subsequently binds fMet-tRNA, the initiator tRNA charged with formylmethionine, to form the 30S initiation complex (30SIC). Dissociation of IF3 yields the intermediate 30SIC’, after which joining of the 50S ribosomal subunit forms the elongation-ready 70S initiation complex.

##### Elongation stage

Elongation is represented by a repeated cycle of four ribosome–tRNA–mRNA states: **EC(*i*), EC’1(*i*), EC’2(*i*)**, and **EC’3(*i*)**. The cycle begins when the ternary complex (TC), consisting of elongation factor Tu (EF-Tu), GTP, and aminoacyl-tRNA (aa-tRNA), binds the ribosomal A-site. EC(0) is initialized from the fully assembled 70S initiation complex in the first elongation round; each subsequent EC(*i*) state is generated from EC’3(*i*-1) after translocation in the preceding round. TC and elongation factor G (EF-G) associate with elongation complexes through the rate constant *k*_on,EF_ adopted from the framework of Dai et al. [10]. The equations use an effective TC concentration (**TC**_**eff**_) equal to 0.028 of total TC to approximate codon-usage and tRNA-abundance heterogeneity [10].

TC association converts EC(*i*) into EC’1(*i*), and peptide-bond formation then generates EC’2(*i*). EF-G association produces EC’3(*i*) and drives translocation along the mRNA. After translocation, EF-G is released and the ribosome enters the next EC state. The index *i* ranges from 0 to the defined average peptide length (*l*_pep_) and enumerates the represented elongation steps.

To reduce computational complexity, repeated elongation reactions were aggregated rather than represented by a separate set of equations for every step *i* = 0, …, *n*, where 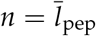. Four equations retain the initial complexes EC(0), EC’1(0), EC’2(0), and EC’3(0). The remaining steps are represented by four lumped variables, defined as 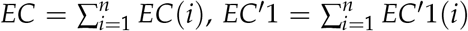, and analogously for EC’2 and EC’3. This aggregation reduces the original 4(*n* + 1) equations to eight differential equations while retaining the repeated association, peptide-bond-formation, and translocation kinetics. It does not explicitly resolve mRNA release according to the ribosomal footprint; instead, RBS release is approximated by the first translocation event from EC’3(0).

GTP hydrolysis during peptide-bond formation releases EF-Tu in its GDP-bound form. EF-Ts-mediated guanine nucleotide exchange (GNEX) regenerates EF-Tu–GTP, which is required for subsequent TC formation. The GNEX module was adapted from the kinetic measurements of Gromadski et al. [18] and their subsequent implementation in the translation model of Hu et al. [22].

##### Termination stage

Termination is represented as three sequential modules: peptide release by RF1 or RF2, release-factor clearance with optional RF3-assisted ribosome rotation, and RRF/EF-G/IF3-mediated ribosome recycling [36, 35]. The corresponding intermediate states distinguish pre- and post-peptide-release complexes, RF-bound rotated states, and the final 30S-containing recycling complexes. This level of detail closes the translation cycle and returns ribosomal subunits for reinitiation; the complete state definitions and ODEs are provided in *Supplementary* Table S1.

#### 4.1.2 Auxiliary modules coupled to translation

Although the analysis centers on initiation, mRNA–ribosome interaction, and ribosome utilization, all simulations used the full coupled translation-cycle model. The auxiliary modules for tRNA charging, EF-Tu/EF-Ts-mediated nucleotide exchange, ribosome stalling, and condition-dependent inhibition of selected GTP-dependent factor-association steps were therefore retained. The last module represents the general inhibitory influence of the (p)ppGpp-mediated stringent response on translational GTPases, including IF2, EF-Tu, EF-G, and RF3 [24].

The model does not simulate ppGpp or pppGpp concentrations, synthesis, or degradation. Instead, normalized static modifiers derived from the RelA/SpoT stoichiometric ratio provide a condition-dependent proxy for inhibition of the implemented IF-dependent progression, TC association, and EF-G association steps. This proxy encodes an effective regulatory trend rather than an alarmone concentration or a resolved model of stringent-response biology. Together, the auxiliary modules keep fMet-tRNA supply, TC availability, free-ribosome recovery, and GTP-dependent association coupled to initiation and ribosome loading. The tRNA-charging module is described in *Supplementary* subsection 1.3.1; the complete ODEs and kinetic constants are listed in *Supplementary* Table S1 and Table S2; and the static proxy is derived in *Supplementary* section 1.9.

#### 4.1.3 Growth-rate-dependent component initialization

For each simulated condition, the input variable was the RNA-to-protein mass ratio, 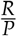, which was mapped to a growth-rate scale using the empirical relationship reported by Dai et al. [10]. Initial molecular abundances were first calculated on a cell-dry-weight basis and then converted into cellular molar concentrations through

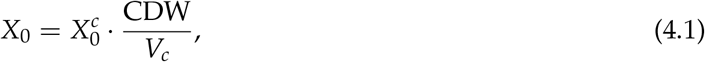

Where 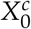 denotes the amount per cell dry weight, *X*_0_ denotes the cellular concentration used by the simulator, and ^CDW^ = 330 g L^−1^ was used following Kremling [27].

The total mRNA amount was estimated from the protein mass fraction, the RNA-to-protein ratio, and the mRNA fraction of total RNA:

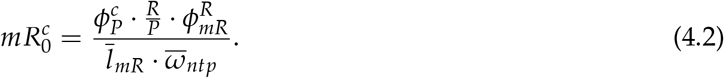

The mRNA fraction 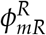 is defined relative to total RNA, rather than relative to cell dry weight. Dai et al. reported a combined tRNA and rRNA fraction of approximately 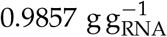, leaving an inferred mRNA fraction of 1 − 0.9857 = 0.0143 g g^−1^ [10]. A factor of 2.3 was then applied to align the reconstructed total-mRNA abundance with the scale reported by Balakrishnan et al., giving the model value 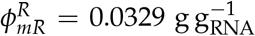 [2]. The corresponding mRNA fraction on the cell-dry-weight scale is therefore the product 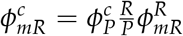, as used in Equation 4.2. The average mRNA length was computed as the coding-sequence contribution plus an untranslated-region contribution of 85 nucleotides [22]:

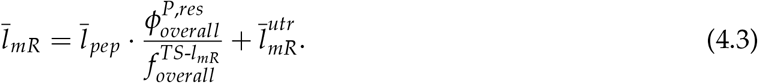

Here, 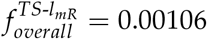 is the transcript-signal contribution normalized by coding-sequence length, and 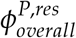 is the peptide-length-normalized proteomic mass contribution. The molecular-weight construction is detailed in *Supplementary* subsection 1.4.1, the mRNA-abundance calibration in *Supplementary* section 1.5, and the omics processing in *Supplementary* section 1.6. The proteome-weighted average peptide length was represented by

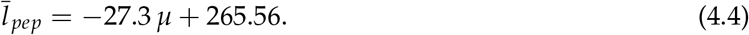

Because this regression has weak explanatory power, it was used only as a coarse-grained initialization term.

The remaining RNA mass after subtracting mRNA was partitioned into tRNA and rRNA using the growth-dependent tRNA-to-rRNA mass ratio adopted from Hu et al. [21]. The resulting 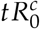 and 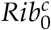 values were converted into molar concentrations using the average tRNA molecular weight and the molecular weight of the complete rRNA complement of one ribosome, respectively. The calculated *Rib*_0_ defines equal initial pools of 30S and 50S subunits. The allocation equations and the growth-dependent tRNA-to-rRNA ratio are provided in *Supplementary* subsection 1.4.2 and Figure S13.

Protein factors and enzymes were initialized as stoichiometric ratios to the ribosome amount:

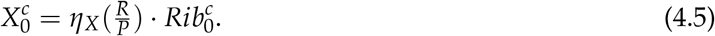

For factors with clear growth-dependent trends, *η*_*X*_ was fitted as a function of 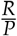 . For IF1, IF2, and RF3, the proteomics-derived trends were weak or inconsistent across datasets, and mean factor-to-ribosome ratios were used. These growth-dependent input functions are treated as coarse-grained initialization and comparison functions rather than as independent biological growth laws. Their derivation, fitted curves, and coefficients are reported in *Supplementary* subsection 1.4.3, Figure S14, and Table S4.

Small-molecule inputs required for nucleotide exchange and tRNA charging were assigned separately. GTP, GDP, and methionine were held constant across the simulated growth range, whereas the total amino-acid concentration followed a prescribed sigmoidal function of 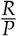. The parameter values, input function, and resulting concentration profiles are provided in *Supplementary* subsection 1.4.4 and Figure S17.

#### 4.1.4 Numerical integration and simulated readouts

The simulation workflow is summarized in Figure 4.2. Inputs included the RNA-to-protein mass ratio 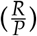, factor-to-ribosome ratios, kinetic rate constants, and the average peptide length 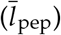. For each 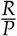 value between 0.09 and 0.55 g_RNA_ g_protein_^−1^, the molecular species were initialized, kinetic parameters were assigned, and the stiff ODE system was integrated from *t* = 0 to 300 s with an implicit Radau solver. Each integration interval contained 200 evaluation points, with relative and absolute tolerances of 10^−8^ and 10^−10^, respectively.

**Figure 4.2.**
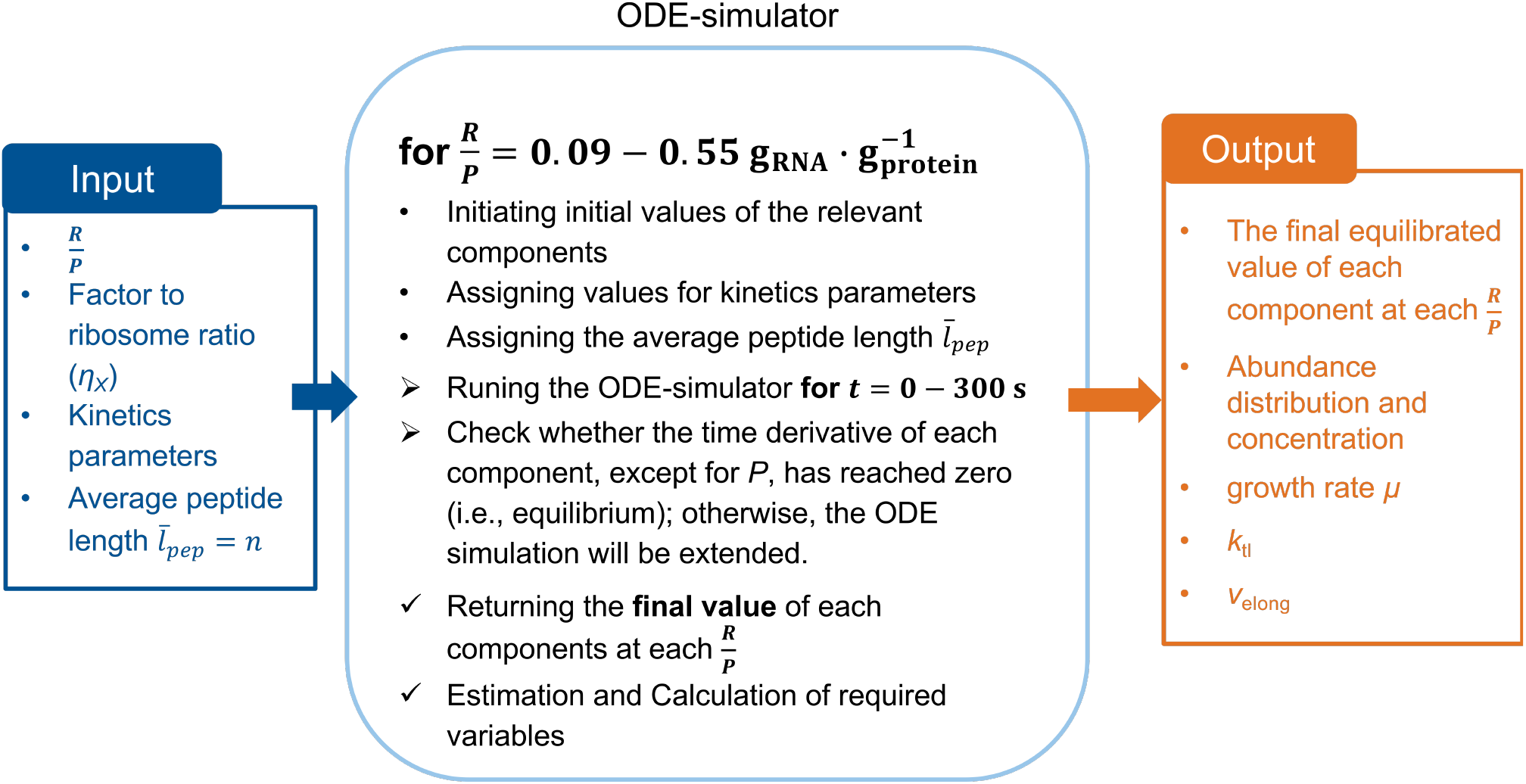
Workflow used to initialize, integrate, extend, and analyze the ODE-based translation simulator across the RNA-to-protein ratio sweep.

Based on the empirical 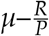 relationship reported by Dai et al. [10], this sweep corresponded to a growth-rate coordinate of approximately *µ* = 0.00–2.11 h^−1^. The full grid was retained in the plotted trajectories to show numerical continuity near the lower boundary. Quantitative endpoint summaries in the Results were restricted to *µ* = 0.10–2.11 h^−1^ because these analyses assume balanced exponential growth and the fixed protein-degradation term strongly influences net protein production near zero growth. The lower reporting limit is therefore a conservative analysis boundary rather than a biological transition point.

After each integration, the final time window was checked for quasi-steady-state convergence of all molecular species except the protein compartment *P*. This exception was necessary because *P* is the accumulated translation product and was used to compute production flux rather than to define molecular-complex equilibrium. If the maximum absolute change across the last five sampled solution intervals exceeded 10^−5^ for any non-protein species, the numerical simulation was extended by another 300 s using the final state from the previous integration as the new initial condition. This extension was repeated up to three times.

After quasi-steady state was reached, the final component concentrations were used to calculate the simulated growth rate (*µ*_simulated_), translation frequency (*k*_tl_), initiation frequency (*k*_ini_), overall elongation velocity (*v*_elong_), and the distributions of ribosomes and translation factors. This workflow links the prescribed macromolecular composition to the resulting translation dynamics.

*Supplementary* section 1.2 and Algorithm 1 provide auditable pseudocode for the parameter sweep, convergence extension, readout extraction, and output storage. The full ODE, parameter, and regression tables are provided in *Supplementary* Table S1, Table S2, and Table S4.

#### Translation and initiation frequencies

The overall translation frequency was estimated by normalizing the net protein production rate by the initial total mRNA abundance:

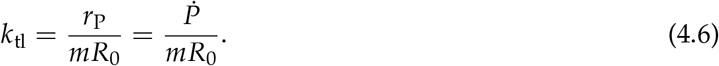

Here, *r*_P_ denotes the net retained protein production rate evaluated after the non-protein molecular species reached quasi-steady state, 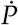 denotes the corresponding simulated protein flux, and *mR*_0_ denotes the initial total mRNA abundance. The flux 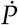 was obtained by numerically differentiating the simulated protein variable *P* with `numpy.gradient`. The same normalization was applied to the COG-specific protein-production and mRNA-template pools defined below.

To recover the nascent synthesis frequency before protein degradation, the modeled degradation contribution was added back to *k*_tl_:

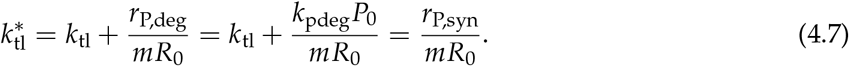

Here, *r*_P,deg_ = *k*_pdeg_*P*_0_ denotes the protein degradation rate, *k*_pdeg_ is the protein-degradation constant, *P*_0_ is the cellular protein concentration for the corresponding condition, and *r*_P,syn_ = *r*_P_ + *r*_P,deg_ is the nascent protein synthesis rate. The value of *k*_pdeg_ was set to 0.03 h^−1^ based on the range reported by Hu et al. [22]. This degradation term is also included in the protein-balance equation (*Supplementary* Table S1). Because the simulator computes a quasi-steady-state distribution under predefined total abundances, *r*_P,deg_ is condition-specific but constant within each 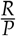 iteration.

Following Kremling’s derivation, the translation initiation frequency is expected to equal 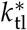 because both describe nascent protein production normalized by mRNA abundance [28]. In the ODE simulator, the initiation frequency (*k*_ini_) was computed independently from the abundance of the last initiation intermediate, 70SPIC, and the kinetic constant of the final initiation step (*k*_i2_):

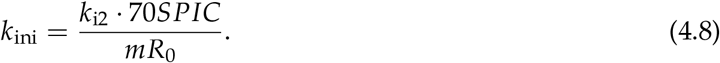

The separately computed *k*_ini_ was compared with 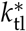 as an internal consistency check.

To calculate COG-specific translation frequencies, gene-level protein and mRNA molar fractions were summed within each COG class to obtain 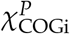 and 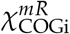, respectively. The class-average translation frequencies were then calculated as

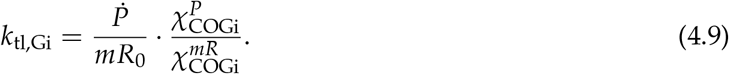

The 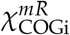 values fluctuate among growth conditions without a consistent growth-rate dependence (Figure 2.1e), whereas 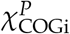 shows clearer class-specific trends (Figure 2.1d). The mean 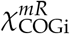 across the available growth conditions was therefore used for each COG class in Equation 4.9. Growth-dependent 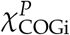 values were obtained from the corresponding linear regressions against *µ*. The simulated growth rate was calculated from the steady-state protein-production rate (*r*_P,*ss*_), the cellular protein mass fraction 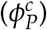, the cell-dry-weight-to-volume ratio 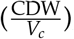, the average amino-acid-residue molecular weight 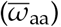, and the average peptide length 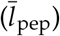 :

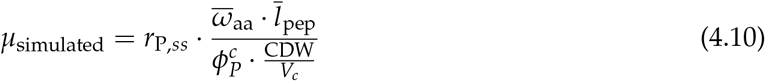

The resulting *µ*_simulated_ was compared with the growth rate reported by Dai et al. at each matched 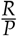 value as a consistency check between the simulated protein-production flux and the empirical growth-rate scale.

#### Elongation velocity and occupancy metrics

The elongation velocity *v*_elong_ was derived from the final quasi-steady-state concentrations of elongation factor G (EF-G) and ternary complex (TC) [10]:

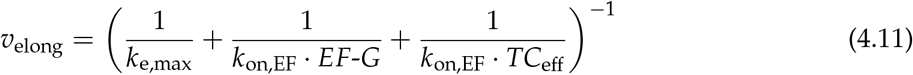

where *k*_on,EF_ is the association rate constant for EF-G and TC, and *k*_e,max_ is the maximal elongation velocity, set to 29 s^−1^. The inverse-sum formulation accounts for the kinetic contributions of maximal translocation capacity, EF-G availability, and effective ternary-complex availability. In the implemented model, the EF-G and TC association terms are multiplied by the condition-dependent static modifiers defined below.

The association steps of EF-G and TC are treated as ribosome-dwelling steps. By contrast, peptide-bond formation and forward elongation are treated as translocation-associated steps with first-order rate constants *k*_pep_ and *k*_esf_. The effective maximal elongation constant is therefore:

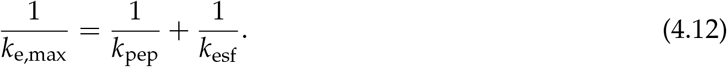

With *k*_pep_ = 60 s^−1^ and *k*_e,max_ = 29 s^−1^, *k*_esf_ was calculated as 56.13 s^−1^.

Because total TC abundance is not equivalent to codon-specific effective TC availability, the effective TC concentration was approximated using the codon-class correction introduced by Dai et al.:

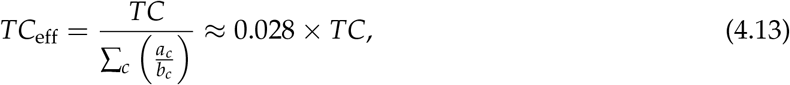

With

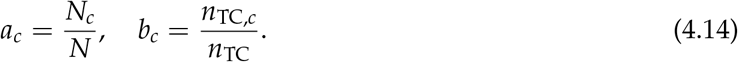

Here, *a*_*c*_ represents the codon abundance fraction of codon class *c*, and *b*_*c*_ represents the corresponding TC abundance fraction. The total translation-associated ribosome pool, *C*_tot_, was calculated as the sum of the initiation, elongation, and termination complexes. For comparison with the amino-acid-carrying active-ribosome fraction reported by Dai et al., the early 30S–mRNA states 30*SRBS* and 30*SPIC*, together with all post-elongation states after peptide releasing (starting from *PoTerCI*1 and *PoTerCI*2 illustrated in Figure 4.1), were excluded from *C*_tot_ before normalization by the initial ribosome amount. Complexes were also grouped by initiation, elongation, and termination stages to quantify phase-resolved ribosome occupancy.

### 4.2 Omics-derived reference estimates and simulation-output analysis

#### 4.2.1 Pool-based reference estimates of average translation frequency

For the main analysis, average translation frequency was estimated at the level of the total expressed protein pool or a COG-specific protein–mRNA pool. This choice reflects the coarse-grained objective of comparing sector-level translation demand and mRNA–ribosome utilization across growth conditions rather than inferring gene-specific translation rates. Gene-wise distribution diagnostics are therefore reported only as *Supplementary* statistical diagnostics.

Under balanced growth, the protein synthesis rate required to maintain a protein pool is approximated by the dilution term *µ* · *P*. The corresponding pool-average translation frequency is:

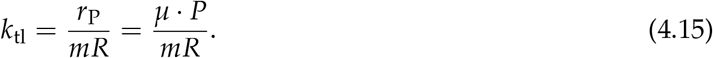

Here, *P* and *mR* denote protein and mRNA abundances expressed on the same physiological scale. For a COG group *i*, the pool method estimates the average translation frequency by aggregating all proteins and mRNAs assigned to that group before taking the ratio:

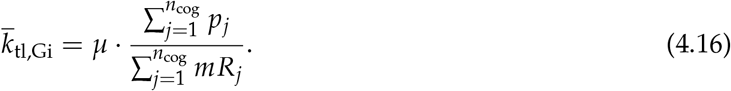

In this formulation, the COG group is treated as a single production pool. This avoids over-weighting low-abundance transcripts and makes the estimated 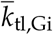 directly comparable with the COG-level protein-production terms used in the ODE simulator.

For the matched proteomics–transcriptomics dataset of Balakrishnan et al. [2], the reported molar fractions of gene-specific protein and mRNA were converted into absolute abundances using the total cellular protein amount *n*_*P*_ and total cellular mRNA amount *n*_*mR*_:

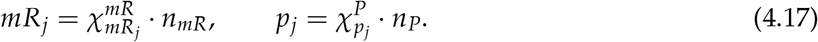

The resulting *p*_*j*_ and *mR*_*j*_ values were then substituted into Equation 4.16. These estimates are used as matched-omics references for the simulated translation-frequency trajectories. The COG mapping and gene-level proteomic and transcriptomic processing are documented in *Supplementary* section 1.6; the alternative gene-wise log-AK calculation is retained as a diagnostic in *Supplementary* section 1.7.

#### 4.2.2 mRNA ribosome-loading fraction

The average number of ribosomes bound to a translated mRNA was estimated from the degradation-corrected nascent translation frequency 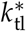, the average elongation velocity *v*_elong_, and the average peptide length 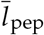. Since 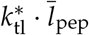 represents the amino-acid polymerization demand per mRNA and *v*_elong_ represents the amino-acid polymerization rate per ribosome, their ratio gives the mean ribosome load per translated mRNA [28, 27]:

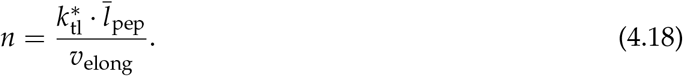

The theoretical maximal ribosome-loading capacity of an mRNA was estimated from the ribosomal footprint, represented by the occlusion length *ℓ*:

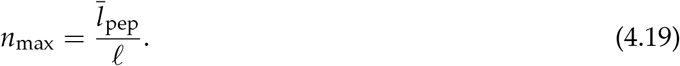

Here, *ℓ* was set to 10 amino acids, consistent with values commonly used to estimate ribosome footprint-limited loading capacity [31, 41, 20, 45]. The fraction of this theoretical capacity that is occupied was then defined as:

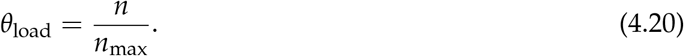

The dimensionless quantity *θ*_load_ is referred to as the mRNA ribosome-loading fraction. It estimates the transcript-wide mean ribosome load as a fraction of the theoretical footprint-limited maximum and was used to evaluate whether the simulated system approaches TX-LIM, where mRNA templates would be densely loaded with ribosomes. It is distinct from the bound-RBS fraction: bound-RBS reports the fraction of mRNA initiation sites represented in mRNA-containing ribosomal complexes, whereas *θ*_load_ estimates ribosome occupancy across the translated coding region. Because the growth-rate-dependent proteome-weighted peptide length is estimated from a weak linear trend, the resulting loading fractions were interpreted as coarse-grained consistency patterns rather than as precise measurements of polysome occupancy.

#### 4.2.3 Descriptive fitting of limitation-regime relationships

The production-rate relationships used to interpret CF-LIM and TL-LIM were fitted to the steady-state simulation outputs. For the relationship between protein production rate, *r*_P_, and free RBS concentration, *RBS*, the points were ordered by *RBS*. Candidate split positions were evaluated with at least six points in each segment. At each candidate position, the low-*RBS* segment was fitted by nonlinear least squares to a Michaelis–Menten function, *r*_P_ = *V*_max_*RBS*/(*K* + *RBS*), and the high-*RBS* segment was fitted by ordinary least squares to *r*_P_ = *aRBS* + *b*. The split that minimized the combined residual sum of squares of the two segments was retained. No continuity constraint was imposed between the two descriptive fits.

Two additional fits were performed over the complete simulated growth range. The dependence of *r*_P_ on total ribosome concentration, *Rib*_0_, was fitted by ordinary least squares, whereas the ribosome-normalized production rate was fitted to *r*_P_/*Rib*_0_ = *V*_max_*RBS*/(*K*_CF_ + *RBS*) by nonlinear least squares with non-negative parameters. Coefficients of determination were calculated against the simulation outputs. These analyses summarize the shape of one continuous growth-dependent trajectory; they were not used to set ODE parameters, calibrate the simulator, or impose a transition between discrete kinetic states.

### 4.3 Kinetic parameter assignment

Kinetic constants were assigned from literature measurements where direct measurements were available. These include initiation steps involving fMet-tRNA binding and 50S joining, elongation-rate constraints, and termination/recycling constants measured in single-molecule or biochemical studies [1, 17, 10, 36, 35]. Because many of these values were measured under *in vitro* conditions, and because several association constants are not available for the exact coarse-grained species used in the simulator, the implemented parameter set combines literature-derived constants, diffusion-theory estimates, temperature-corrected estimates, and calibrated effective constants. The full parameter table is provided in the *Supplementary Information* (Table S2).

For association steps lacking directly usable *in vivo* rates, pair-specific diffusion-limit scale estimates were calculated using

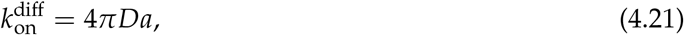

where *D* = *D*_*A*_ + *D*_*B*_ is the relative diffusion coefficient of the interacting molecules and 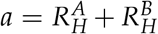 is the encounter distance. Hydrodynamic radii were estimated from molecular weight using

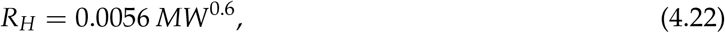

where *R*_*H*_ is in nm and *MW* is in Da. To avoid representing the cytoplasm as a uniform Newtonian medium, the Stokes–Einstein calculation assigned each component an effective viscosity determined by its hydrodynamic radius using the *E. coli* scale-dependent viscosity reference curve [25]. The two component-specific diffusion coefficients were then summed; their effective viscosities were not averaged. A second calculation using water viscosity at 37 ^*°*^C provided an aqueous reference scale. Association constants were converted from µm^3^ molecule^−1^ s^−1^ to µM^−1^ s^−1^ using 1 µm^3^ molecule^−1^ ≈ 602 µM^−1^. The resulting cytoplasmic and aqueous estimates are compared with the implemented association constants in *Supplementary* Table S3; the effective-viscosity calculation and unit conversion are detailed in *Supplementary* subsection 1.8.1. These values were used as physically motivated comparison scales, not as measured intracellular microscopic constants or direct replacements for literature-derived and calibrated parameters.

Termination constants measured at temperatures below 37 ^*°*^C were adjusted where possible using an Arrhenius relationship between ln *k* and 1/*T*. When only one member of a mechanistically related pair had enough temperature information, the same temperature-scaling ratio was applied to the corresponding paired rate. These corrections were used only to place termination kinetics on a plausible 37 ^*°*^C scale; the rate-pair assignments and calculations are documented in *Supplementary* subsection 1.8.2.

Three initiation-side quantities were treated as calibrated effective parameters. The parameter *k*_a,30S_ describes effective 30S–mRNA/RBS association, and *k*_IF_ describes a lumped IF-dependent progression step that multiplies the early 30S–mRNA complex by the free concentrations of IF1, IF2, and IF3. The third parameter, ℋ_I,IF_, controls the shape of the normalized static IF-related modifier described below. None of these three quantities corresponds to a single microscopic elementary reaction. In the simulator’s mM concentration scale, the baseline values are *k*_a,30S_ = 0.7 *×* 10^3^ mM^−1^s^−1^, *k*_IF_ = 1.0 *×* 10^7^ mM^−3^s^−1^, and ℋ_I,IF_ = 1. The *k*_a,30S_ value is equivalent to 0.7 µM^−1^s^−1^. These values were manually adjusted to reproduce the growth-rate scale defined by the empirical 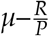 relationship and the active-ribosome-fraction trend reported by Dai et al. [10]. They should therefore be interpreted as calibrated effective parameters rather than as direct literature measurements.

### 4.4 Static representation of stringent-response-associated inhibition

During the stringent response, (p)ppGpp can inhibit several translational GTPases. Reported targets include IF2 during initiation, EF-Tu-mediated aminoacyl-tRNA delivery, EF-G-mediated translocation, and RF3-mediated recycling of RF1 and RF2 [24]. This established inhibitory effect motivates inclusion of a condition-dependent regulatory term in the simulator; the representation used here is an effective modeling approximation rather than a reconstruction of stringent-response signaling.

The model does not explicitly simulate ppGpp or pppGpp concentrations or their time-dependent synthesis and degradation. Instead, the condition dependence is represented by normalized static modifiers derived from the RelA/SpoT stoichiometric ratio, 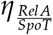. For each implemented step *j*, the effective kinetic rate is

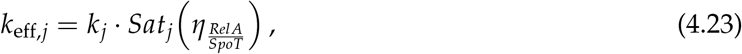

where *Sat*_*j*_ is a normalized saturation term. The modifier is applied to the lumped IF-dependent progression step, TC association during elongation, and EF-G-associated translocation; RF3 inhibition is not implemented. Because IF1, IF2, and IF3 are represented as coordinated participants in the lumped initiation step, the IF-related modifier should not be interpreted as a separately resolved molecular interaction for each factor.

The RelA/SpoT ratio is used only as a static proxy and is not interpreted as a simulated alarmone concentration. The step-specific functions were normalized on a fixed physiological reference grid and calibrated against the growth-rate scale, elongation-velocity trend, and active-ribosome-fraction reference reported by Dai et al. [10]. These measurements are calibration constraints, not independent validation of the proxy or evidence that it reproduces intracellular stringent-response dynamics. The monotonic proxy derivation is provided in *Supplementary* subsection 1.9.2, and the normalization and step-specific functions are given in *Supplementary* subsection 1.9.3.

### 4.5 Calibration and validation status

The model combines empirical initialization rules, calibrated effective parameters, literature-based kinetic estimates, and output-level consistency checks. To avoid treating calibration constraints as independent validation, Table 4.1 separates the main evidence sources by their role in the workflow. Items marked as calibration constraints were used to set model scale or tune effective parameters and were therefore not interpreted as independent validation tests. The corresponding calibration and output-level consistency plots are collected in *Supplementary* section 1.10.

**Table 4.1.** Calibration and validation status of major evidence sources used in the translation simulator.

| Evidence source or check | Use in the manuscript | Workflow role | Independent validation status |
| --- | --- | --- | --- |
| Dai et al. $\mu-\frac{R}{P}$ relationship [10] | Maps the empirical growth-rate coordinate to RNA-to-protein ratio and initializes RNA and ribosome pools. | Initialization input | No; defines the simulated growth coordinate. |
| Dai et al. active-ribosome fraction and elongation trends [10] | Set the active-ribosome scale and constrain selected effective kinetic modifiers. | Calibration constraint | No; used for calibration consistency. |
| Balakrishnan et al. matched proteome-transcriptome data [2] | Provides pool-based translation-frequency references for overall and COG-level trends. | Consistency reference | Limited; dataset- and strain-specific comparison. |
| Harmonized mRNA-abundance estimates from Valgepea et al. and Balakrishnan et al. [46, 2] | Provide the total-mRNA scale after conversion to consistent concentration units. | Consistency reference | Limited; depends on conversion and mass-fraction calibration. |
| Public reproduction code and tests | Audits implementation structure, baseline-grid invariance, and selected numerical outputs. | Reproducibility check | No; supports computational reproducibility, not biological validation. |

### 4.6 Use of AI-assisted tools

OpenAI Codex was used to assist with manuscript organization, language editing, LaTeX formatting, internal consistency checks, and drafting of auxiliary reproduction-code and visualization-notebook text. AI-assisted outputs were reviewed, edited, and verified by the authors against the implemented model, source data, and cited literature. Model assumptions, parameter choices, scientific interpretations, and the final manuscript content were determined and approved by the authors, who take full responsibility for the article.

## Data availability

The study uses published physiological, proteomic, and transcriptomic datasets, which are cited in the main text and the *Supplementary Information*. The ODE simulator, model inputs, generated simulation outputs, and processed data underlying the simulation-based figures and most Supplementary figures are archived together with the reproduction scripts and analysis notebooks at Zenodo: https://doi.org/10.5281/zenodo.22086884. Additional source data and analysis files underlying Figures 2.1d and 2.1e and Supplementary Figures S9–S12 are provided separately with this submission.

## Code availability

The ODE simulator, reproduction scripts, and analysis notebooks are archived at Zenodo: https://doi.org/10.5281/zenodo.22086884.

## Acknowledgements

This work was supported by the Deutsche Forschungsgemeinschaft (DFG, German Research Foundation) under grant KR 2963/9-1.

## Author contributions

J.Q. developed the model implementation, processed the data, performed simulations, generated figures, and drafted the manuscript. A.K. supervised the study, contributed to model conceptualization, and revised the manuscript. All authors reviewed and approved the manuscript.

